# Long-term ecosystem nitrogen limitation from foliar δ^15^N data and a land surface model

**DOI:** 10.1101/2021.07.16.452605

**Authors:** Silvia Caldararu, Tea Thum, Lin Yu, Melanie Kern, Richard Nair, Sönke Zaehle

## Abstract

The effect of nutrient availability on plant growth and the terrestrial carbon sink under climate change and elevated CO_2_ remains one of the main uncertainties of the terrestrial carbon cycle. This is partially due to the difficulty of assessing nutrient limitation at large scales over long periods of time. Consistent declines in leaf nitrogen (N) content and leaf δ^15^N have been used to suggest that nitrogen limitation has increased in recent decades, most likely due to the concurrent increase in atmospheric CO_2_. However, such datasets are often not straightforward to interpret due to the complex factors that contribute to the spatial and temporal variation in leaf N and isotope concentration. We use the land surface model QUINCY, which has the unique capacity to represent N isotopic processes, in conjunction with two large datasets of foliar N and N isotope content. We run the model with different scenarios to test whether foliar δ^15^N isotopic data can be used to infer large scale nitrogen limitation and if the observed trends are caused by increasing atmospheric CO_2_, changes in climate or changes in sources of anthropogenic N deposition. We show that while the model can capture the observed change in leaf N content and predicts widespread increases in N limitation, it does not capture the pronounced, but very spatially heterogeneous, decrease in foliar δ^15^N observed in the data across the globe. The addition of an observed temporal trend in isotopic composition of N deposition leads to a more pronounced decrease in simulated leaf δ^15^N. Our results show that leaf δ^15^N observations should not, on their own, be used to assess global scale N limitation and that using such a dataset in conjunction with a land surface model can reveal the drivers behind the observed patterns.

## 1 Introduction

The magnitude and dynamics of the terrestrial carbon sink remains one of the largest uncertainties in predictions of future climate change (Jones & Friedlingstein, 2020), specifically in relation to the amount of carbon that can be stored in terrestrial ecosystems under future levels of atmospheric carbon dioxide (CO_2_) (Walker et al., 2020). One particular source of uncertainty is the degree to which nitrogen limits plant growth (Elser et al., 2010; Hungate et al., 2003). Additionally, anthropogenic nitrogen input into ecosystems as atmospheric deposition can alleviate nitrogen limitation (Matson et al., 2002). While the processes relating nutrient availability to individual plant physiology are relatively well known, the ecosystem level limitation is poorly understood partially because it is very difficult to measure (Vicca et al., 2018). We propose to use a land surface model to bridge some of these observational gaps and provide insights into the underlying processes.

A number of metrics for ecosystem nutrient limitation have been suggested, both through soil and vegetation metrics (Van Sundert et al., 2020). While soil metrics provide a more comprehensive view of the system, plant-related measurements are much more prevalent, largely due to the ease of sample collection and measurement when compared to soil observations. To assess N limitation under elevated CO_2_ globally, we need measurements that are spatially distributed but also cover a sufficient time period to detect the effects of increased CO_2_. While organised efforts to collect ecosystem data in such a systematic manner exist (e.g. The International Co-operative Programme on Assessment and Monitoring of Air Pollution Effects on Forests (ICP Forests, Lorenz, 1995), their extent is limited both in space and time, and the exact focus of the measurement varies. Thus To have a global picture we must rely on measurements that are general and simple enough to have been made at a large number of studies.

Leaf nitrogen (N) content has been proposed as a diagnostic of ecosystem N limitation, with low leaf N considered a sign of poor plant nutrition (Elser et al., 2010). Generally, controlled experiments show N additions lead to an increase in foliar N (Magill et al., 2004; Sikström, 2002), while elevated CO_2_, generally leads to a decrease (Ellsworth et al., 2004). Based on this metric, previous studies have concluded that forest nutrition is degrading concurrently with increased atmospheric CO_2_ (Jonard et al., 2015). However, plants can alter their leaf chemistry under elevated CO_2_ in various ways, including plastically reducing the amount of N allocated to photosynthetic components due to increased nitrogen use efficiency under elevated CO_2_ (Ainsworth & Long, 2005; Drake et al., 1997). This allows plants to maintain similar levels of biomass growth with less available N. These observations raise the question of whether a decrease in leaf N indicates N limitation to growth or just plant adaptive response to N availability, with little or no effect on growth.

Another proposed metric of N limitation is foliar δ^15^N. Nitrogen isotopic concentrations are a useful tool in assessing ecosystem nitrogen related processes, as processes fractionate differently (Robinson, 2001). In particular, the balance between microbial processes that fractionate and physical processes such as leaching that do not can give an indication of the amount of mineral N available in the system (Amundson et al., 2003). While still a plant-based metric, it can reflect whole-ecosystem processes, with a change in δ^15^N indicating a change in N loss pathways from the system, with less N lost through leaching processes when there is overall less mineral N present in the soil (Amundson et al., 2003; Craine et al., 2015; Hogberg, 1998). This decrease in δ^15^N with decreased N availability is observed in both fertilization experiments (Choi et al., 2005; Johannisson & Högberg, 1994) and across fertility gradients (Garten & Van Miegroet, 1994), although such studies do not generally cover a large spectrum of ecosystems and plant functional types. A recent synthesis of δ^15^N measurements (Craine et al., 2009, 2018) concluded that foliar δ^15^N has decreased over the last decades and this decrease is caused by N limitation, concurrent with the trend in foliar N concentration from long-term monitoring of European forests (Jonard et al. 2015).The study advanced the hypothesis that inferred N limitation is driven by elevated CO_2_

A complication in attribution of trends in foliar N and its isotopic composition to elevated CO2 is the alteration of the N cycle due to anthropogenic N deposition (Galloway et al., 2004). Nitrogen deposition is spatially and temporally complex, and regions of the world such as Europe which before 1990 suffered from chronic high level of N inputs in acid deposition have implemented legislation to limit emissions, while other parts of the world have increased emissions (and thus, deposition) through industrialisation and intensification of agriculture (Fowler et al., 2013; Vitousek et al., 1997). Both fossil fuels and agricultural emissions also have a depleted ^15^N content compared to most ecosystem pools, which can potentially lead to a decreasing ecosystem δ^15^N trend throughout the 20th century due to chronic inputs (Hastings et al., 2009). Additionally the balance of emissions with different δ^15^N have changed over time, with a decrease in NO_x_ from fossil fuels and increase from agricultural emissions leading to a change in δ^15^N of deposition (Elliott et al., 2019; Felix et al., 2012). While exhaustive data of trends in N deposition δ^15^N are not available, large scale trends have been observed in lake sediments (Holtgrieve et al., 2011) and ice cores (Hastings et al., 2009), indicating these effects are significant and detectable.

Land surface models (LSMs) provide detailed representations of ecosystem and landscape level processes and have over the last decade advanced from carbon-only representations to having fully integrated nutrient cycles (Arora et al., 2020; Davies-Barnard et al., 2020). This advancement in process representation means such models can not only be used for prediction, but also interrogating quantities and processes that are difficult to measure. Models can also be used to perform virtual experiments, varying the climate or CO2 concentrations, to further investigate the drivers of observed patterns. This is particularly useful when looking at leaf δ^15^N observations, given the large number of processes involved in N isotope discrimination within an ecosystem, and relatively small changes over time make process attribution difficult. However, the majority of models do not have the capacity to represent N isotopic processes. Here, we use the QUINCY model (QUantifying Interactions between terrestrial Nutrient CYcles and the climate system), a model specifically designed to incorporate a detailed and accurate representation of nutrient cycling processes (Caldararu et al., 2020; Thum et al., 2019). In addition to the common nitrogen cycle components, QUINCY includes nitrogen isotope fractionation processes, which allows us to use the model alongside δ^15^N observations to investigate changes in nutrient limitation.

We use the QUINCY model, in conjunction with the foliar δ^15^N dataset of (Craine et al., 2018) and a European database of leaf N concentrations to test the following hypotheses:

1. Increased atmospheric CO2 is the main driver for the observed trend in foliar δ^15^N and N content
2. Changes in the δ^15^N composition of anthropogenic N deposition contribute to the observed δ^15^N trend.

## 2 Methods

### 2.1 Model description

In this study, we use the land surface model, QUINCY (QUantifying Interactions between terrestrial Nutrient CYcles and the climate system), which includes fully coupled carbon, nutrient (nitrogen and phosphorus), water and energy cycles as well as detailed biological representations of plant and soil processes. The model is described in detail in Thum et al. (2019) and here we provide a brief description of the model, with more detailed descriptions of the processes relevant to this study following below.

QUINCY includes a multi-layer canopy representation, with photosynthesis and stomatal conductance being calculated separately for sunlit and shaded leaves for each layer. Nitrogen is vertically distributed through the canopy, with exponentially decreasing N content towards the bottom-most layer. Photosynthetic N is used to calculate photosynthesis for each canopy layer, using the model of Kull & Kruijt (1998). Plant maintenance respiration is calculated as a linear function of N content for each vegetation pool. Both photosynthesis and respiration parameters are acclimated to temperature following Friend (2010) and Atkin et al. (2015) respectively. Plant nutrient uptake is calculated as a function of fine root biomass, soil mineral nutrient concentration and plant nutrient demand.

In QUINCY, sink and source processes are separated through the introduction of two non-structural biomass pools, the short-term labile and the long-term reserve pool. Newly acquired C, N and P are allocated to the labile pool, from where they are further allocated to new growth, respiration or storage in the reserve pool. Actual growth is determined by the nutrient constraint set by the stoichiometry of each plant pool, the nutrients available for growth and further controlled by air temperature and soil moisture. We represent plant nutrient use efficiency response to nutrient limitation through changes in tissue stoichiometry and allocation shifts between foliage and fine roots. Plant N content varies following the optimality scheme described in Caldararu et al. (2020) in which plants adjust their N content in order to maximise growth given current environmental conditions and nutrient availability. Leaf to root ratio is calculated following an empirical function which is based on the ratio between the plant nutrient demand for growth and the available nutrients. In terms of vegetation turnover, QUINCY includes density dependent mortality and establishment from a seed-bed pool. All pools and fluxes are representative of an average individual.

The soil component of QUINCY has a vertically layered structure (with 15 layers of exponentially increasing depth up to a total soil depth of 9.5 m), with processes largely following the CENTURY model approach (Parton et al., 1993). Each soil layer contains five organic pools (metabolic, structural and woody litter, fast and slow overturning soil organic matter (SOM)), as well as inorganic N (NH4+ and NO3-). All litter and SOM pool turnover follows first-order kinetics functions, with soil temperature and moisture dependencies. Litter stoichiometry is determined by the respective plant pool stoichiometry and the plant resorption capacity for each nutrient, while the stoichiometry of the fast SOM pools depends on available inorganic nutrients and the slow SOM pool has fixed stoichiometry. Plant uptake and microbial immobilisation (SOM decomposition) compete for inorganic nutrients based on their uptake capacity and nutrient demand. QUINCY includes a representation of nitrification and denitrification processes based on the aerobic status of each soil layer, resulting in the emission of NO_y_ and N_2_O respectively (Zaehle & Dalmonech, 2011). Leaching of mineral nutrients from the system is represented as a function of mass water flow and nutrient concentration. Biological nitrogen fixation (BNF) is considered as an asymbiotic as well as symbiotic process and is calculated based on plant demand and relative costs of root uptake of mineral nutrients and biological fixation (Meyerholt et al., 2016; Rastetter et al., 2001). Atmospheric nitrogen deposition is a model input (Section 2.5.1) and is added as ammonia and nitrate to each of the respective mineral soil pools in the top layer at every timestep.

While QUINCY has the capacity to represent both N and P processes, for the purpose of this analysis we run the model with P availability set to a level at which is does not limit plant and soil processes, and plant and soil P stoichiometry represent average conditions based on literature values Thum et al. (2019).

### 2.2 Representation of nitrogen stable isotopes in QUINCY

All soil and vegetation pools in QUINCY contain C, N, as well as their isotopes, ^13^C, ^14^C and ^15^N. Processes that fractionate with respect to the nitrogen isotopes are biological nitrogen fixation, ammonification, plant and microbial N uptake, and processes associated with nitrification and denitrification. All isotopic fractionation is described and parameterized following Robinson (2001, Table S2). We calculate leaf δ^15^N as:

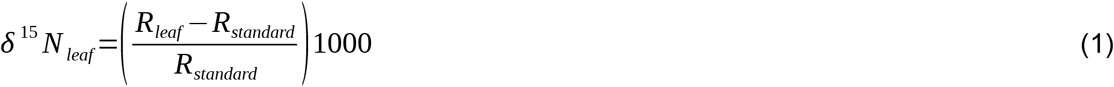

where R_leaf_ is the isotopic ratio of the leaf and R_standard_ is the atmospheric N_2_ standard isotopic ratio (0.0036765).

To showcase the model behaviour in terms of N isotopes, we run the model for one site across a gradient of soil N to test the predicted response of leaf δ^15^N to N availability and increased CO_2_. As expected from theory, the model predicts an increase in leaf δ^15^N with soil N content and a lower leaf δ^15^N under elevated CO_2_ (Fig. 1(a)), reflecting our process understanding of 15N fractionation at the ecosystem level. The ratio between plant N uptake to N loss from the system (Fig. 1(b)) is a measure of the openness of the nitrogen cycle, which can be used as a proxy for N limitation - a low ratio indicates excess mineral N in the system, not taken up by plants and available to gas loss and leaching. Again, as expected from theory, QUINCY predicts a decrease in leaf δ^15^N with an increase in the uptake to loss ratio and therefore with decreased N availability.

**Figure 1.**
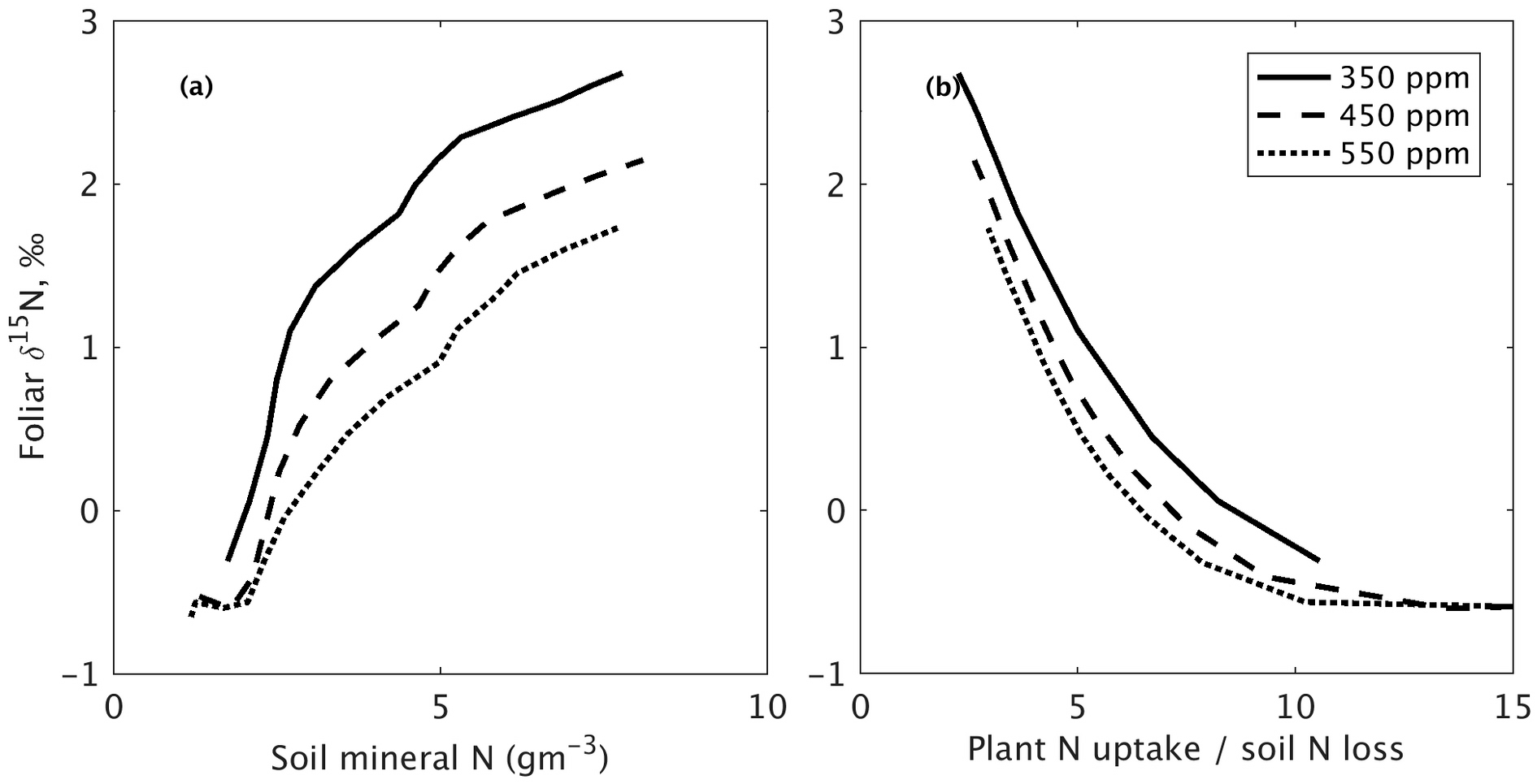
Conceptual model predictions of leaf d15N variation at three levels of atmospheric CO2 for ecosystems with different levels of N availability, plotted against (a) simulated soil N availability and (b) simulated plant N uptake to ecosystem N loss ratio. Simulations shown for a needleleaf evergreen site, and were derived by varying the level of asymbiotic BNF input in the model, which effectively changes the amount of mineral N in the soil. For the purpose of this run only we assume that BNF does not fractionate. This conceptual run removes the effects of climate, PFT and N deposition. We run the model to equilibrium for 500 years with static climate and with constant atmospheric CO2 set to 350 ppm, 450 ppm and 550 ppm in three separate model runs.

### 2.3 Metrics of ecosystem N status

As discussed above, there is no standard metric for assessing nutrient limitation at the ecosystem level. The advantage of using a model in this context is that we can calculate a number of different metrics to describe the N limitation predicted by our model. In this study, we will use:

- **Leaf nitrogen content** - growing season yearly average of total canopy leaf N content. The growing season is defined as periods with GPP (gross primary productivity) above 20% of the maximum at each site.
- **C:N ratio of the plant labile pool** - annual mean of the C:N ratio of material available for plant growth. Higher values indicate a stronger plant N limitation.
- **Soil mineral N** - annual mean of soil soluble NH_4_ and NO_3_ pools to 1 m depth.
- **Ratio of ecosystem uptake to loss** - ratio of plant N uptake to sum of leaching and gas loss fluxes. All represented as annual fluxes. An increase in this ratio shows a more closed N cycle, associated with increasing ecosystem N limitation.

### 2.4 Datasets used

We use two long-term datasets, the foliar ^15^N global data of (Craine et al., 2018) (hereafter Craine2018) and the ICP Forests Europe-wide leaf N content data (Lorenz, 1995). The Craine2018 dataset consists of foliar ^15^N measurements gathered from literature, as well as previously unpublished data. The dataset contains measurement from 1902 to 2017 and covers sites across the globe (Fig. S1(a)) but following the original study, we only use data from 1980 onwards. It contains information on foliar δ^15^N, foliar N content, species, mean annual temperature, mean annual precipitation, mycorrhizal association and nitrogen fixing capacity. For the purpose of this analysis we exclude N-fixing species, as QUINCY does not currently include an N fixer plant functional type. We include additional information on plant functional type based on species information using the categorical traits dataset from the TRY database (Kattge et al., 2020) and N deposition values from the same global dataset as used for the model simulations as described below (Lamarque et al., 2010, 2011).

The International Co-operative Programme on Assessment and Monitoring of Air Pollution Effects on Forests (ICP Forests) is a Europe-level monitoring network with regular standardised measurements of plant, soil and ancillary observations at established forest plots (Fig. S4). Here, we use data for leaf N content from 1990 to 2015.

### 2.5 Model setup

#### 2.5.1 Boundary conditions and meteorological forcing

We run the QUINCY model at approximately 400 sites, distributed uniformly across climate zones and plant functional types (Fig. S1(b), Table S3) for the period 1901-2018. The modeled sites cover the climate space and the range of the observations (Fig. S2). We use daily meteorological data (short and longwave downward radiation, air temperature, precipitation, air pressure, air humidity and wind velocity) extracted from the CRU JRA dataset, version 2.1 (Harris, 2020), and disaggregated to the half-hourly model timestep using a statistical weather generator (Zaehle & Friend, 2010). As other model inputs we use annual atmospheric CO_2_ concentration from Le Quéré et al. (2018) and N deposition data from Lamarque et al. (2010, 2011). QUINCY also requires information about the dominant plant functional type (Hurtt et al., 2006), and soil physical and chemical parameters obtained from the SoilGrids dataset (Hengl et al., 2017)

We bring the model to quasi-equilibrium using a spinup period of 500 years, using driving data from 1901-1930. We then run the model with transient climate, CO_2_ and N deposition for 1901-2018 for each site.

#### 2.5.2 Model scenarios for hypothesis testing

To explore the different factors that can contribute to the observed δ^15^N trend and ecosystem N limitation, we run the model with different factorial combinations of drivers (Table 1). The baseline version includes transient climate, CO_2_ and N deposition. The ‘fixed CO_2_’ model scenario, in contrast, includes transient climate but the atmospheric CO_2_ concentration is set to the value for 1901, to test hypothesis 1 that it is the change in CO_2_ concentration, rather than the change in climate that drives any observed N limitation.

**Table 1.**
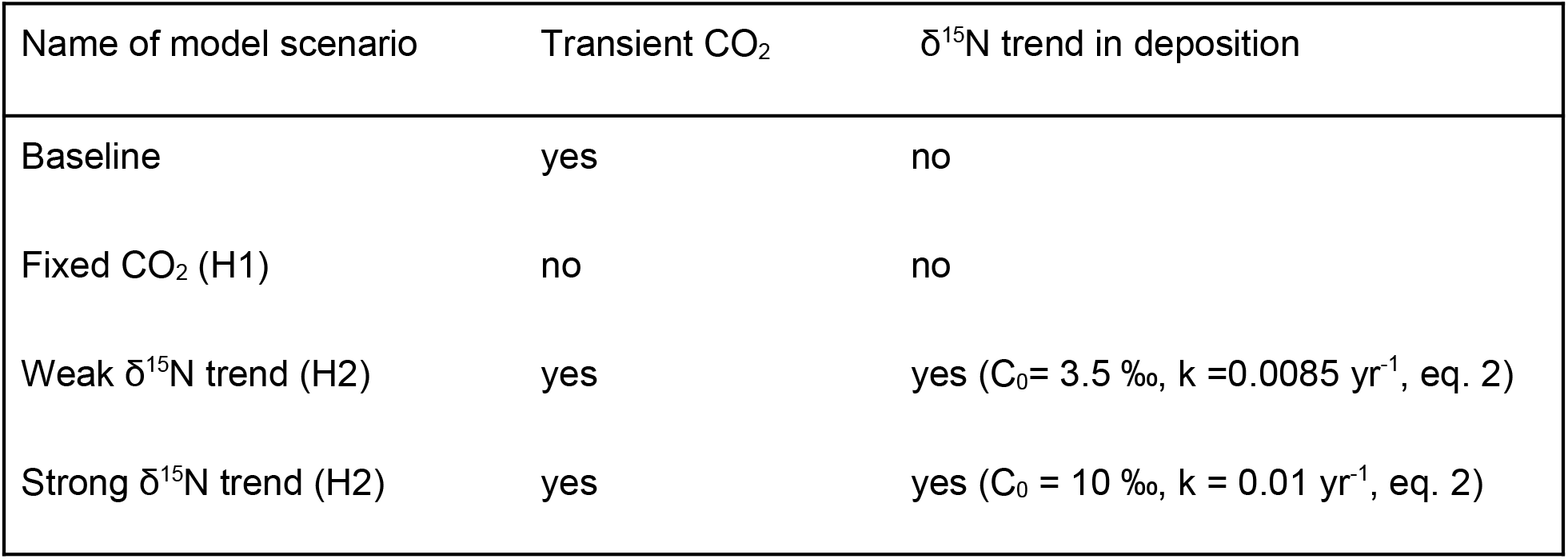
Model scenarios used in this study.

To investigate the effect of a change in the δ^15^N of anthropogenic N deposition, we implement two different scenarios, based on existing observations of long-term trends in lake sediment cores across the US (Holtgrieve et al., 2011) and a Greenland ice core (Hastings et al., 2009). Both records show a decrease in δ^15^N with time over the last century, but they differ in the strength of the trend, with the lake sediments showing a weaker trend than the ice core, possibly due to differing sources for atmospheric transport. Lacking further information on the global distribution of such δ^15^N trends, we test the importance of the N deposition effect using both trends, as a proof of concept of the effect that such a trend would have on foliar δ ^15^N. The ‘weak δ^15^N trend’ scenario uses the data from Holtgrieve et al. (2011), while the ‘strong δ^15^N trend’ scenario follows the trend observed in the Greenland ice core (Hastings et al., 2009). We have implemented both trends following the functional form of (Holtgrieve et al., 2011), so that the δ^15^N concentration in deposition is calculated as:

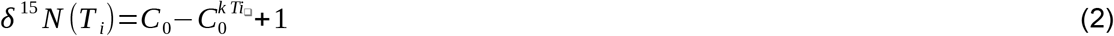

Where *C_0_* is the pre-industrial δ^15^N concentration and *k* is a parameter. The time *T* is expressed as the difference between the simulation year and a threshold year *Y_0_* when the effects of human impact on N deposition had begun, set to 1985 according to the global parameter derived by Holtgrieve et al. (2011). Values for *k* and *C_0_* are taken as the mean of fitted parameters from Holtgrieve et al. (2011) for the weak δ^15^N trend model and are fit to reproduce the ice core timeseries from Hastings et al. (2009) for the strong δ^15^N trend model. Parameter values are included in Table 1.

### 2.6 Statistical analysis

To compare the trends observed in the Craine2018 data and those predicted by QUINCY, we apply a linear regression model to both the observed and predicted values, to account for the effect of spatial variation in mean annual temperature, mean annual precipitation and leaf N content as:

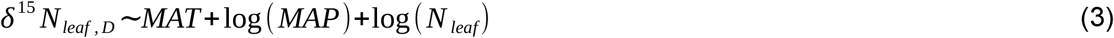

Where *D* denotes either observations or each model scenario. Mean annual temperature (*MAT*) and mean annual precipitation (*MAP*) are climatological means reported in the Craine2018 dataset for the observations and means over the analysis period for the model. Leaf N values are data reported for each site for the observations and means over the analysis period for the model.

We then compute the residuals of this linear regression model for both observations and data, 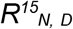. By using the residuals rather than the raw values, we control for spatial variation in the climatic drivers of leaf δ^15^N. We then calculate the trend by computing a linear regression of mean annual residuals with time separately for the observation, each model scenario and each grouping of interest:

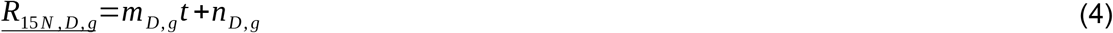

Here *t* is time expressed as year from 1901, *m* and *n* are the slope and intercept respectively, *D* denotes observation or model scenario and *g* is group of interest as either PFT, continent or level of N deposition. For the remainder of this paper, we report the slope *m_D,g_* and its associated standard error. This final regression with time is weighted by the natural logarithm of data points in each year, to account for the high temporal variation in data availability, with more recent years having a larger number of observations (Fig. S3). We report p-values for each slope, as obtained from the linear regression coefficients. Our analysis largely mirrors the one in the original study with the exception of including continent as an explanatory variable in the regression model which we do not do, as we believe that patterns at the continent scale can reflect patterns in N deposition trends and should not be factored out, as we detail in our analysis below. Additionally, mycorrhizal type is not included as a factor in the first regression model (Eq. 3) as this information is not available in the model. This statistical analysis is necessary due to the fact that this dataset does not include continuous timeseries at any one site and thus we must control for climate factors that affect inter-site variability. All statistical analyses were performed using Matlab v9.1 (R2016b) standard functions.

As the ICP Forests data provides timeseries for continuous monitoring plots, we perform a linear regression at each site with time as:

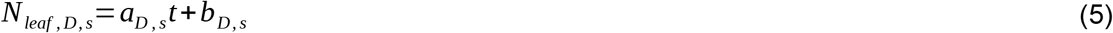

Where *N_leaf,D,s_* is the leaf N content at each site s for either the ICP Forests observation or each model scenario (*D*) and *a* and *b* are the slope and intercept respectively. We report fractional changes per year relative to leaf N values in 1990 calculated as:

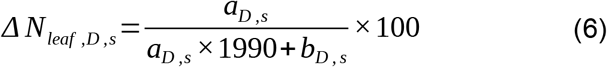

For each PFT we then report the mean of that PFT and the standard error across that PFT. This applies also to the predicted N status metrics described above.

## 3 Results

### 3.1 Global patterns of foliar δ^15^N variation

We explore the variation of δ^15^N residuals across the global gradient with mean annual precipitation, temperature and leaf N content for both model and data.The model shows similar patterns to those observed in the data, with δ^15^N residuals increasing with mean annual temperature and leaf N content, and decreasing with mean annual precipitation (Fig. 2). The slopes of the relationships are generally comparable between the data and model, with a lower model slope for the leaf N content relationship predicted by the model. This discrepancy between data and model for the relationship with leaf N is due to the fact that leaf N content is closely linked to plant function and vegetation type and its relationship with ^15^N content is therefore affected more than that of the climate variables by the differences in site distribution across PFTs between data and model. The few high modeled δ^15^N values (Fig. 2(d) and (e)) are dry and warm sites which, while the climate space of the modeled and observed sites are similar, are not represented in the observed sites. These general patterns indicate that our implementation of ^15^N processes can represent observed global patterns. These results provide confidence that the model can simulate observed patterns in ^15^N across large scales and climatic gradients and therefore supports our following analysis of temporal trends in leaf δ^15^N.

**Figure 2.**
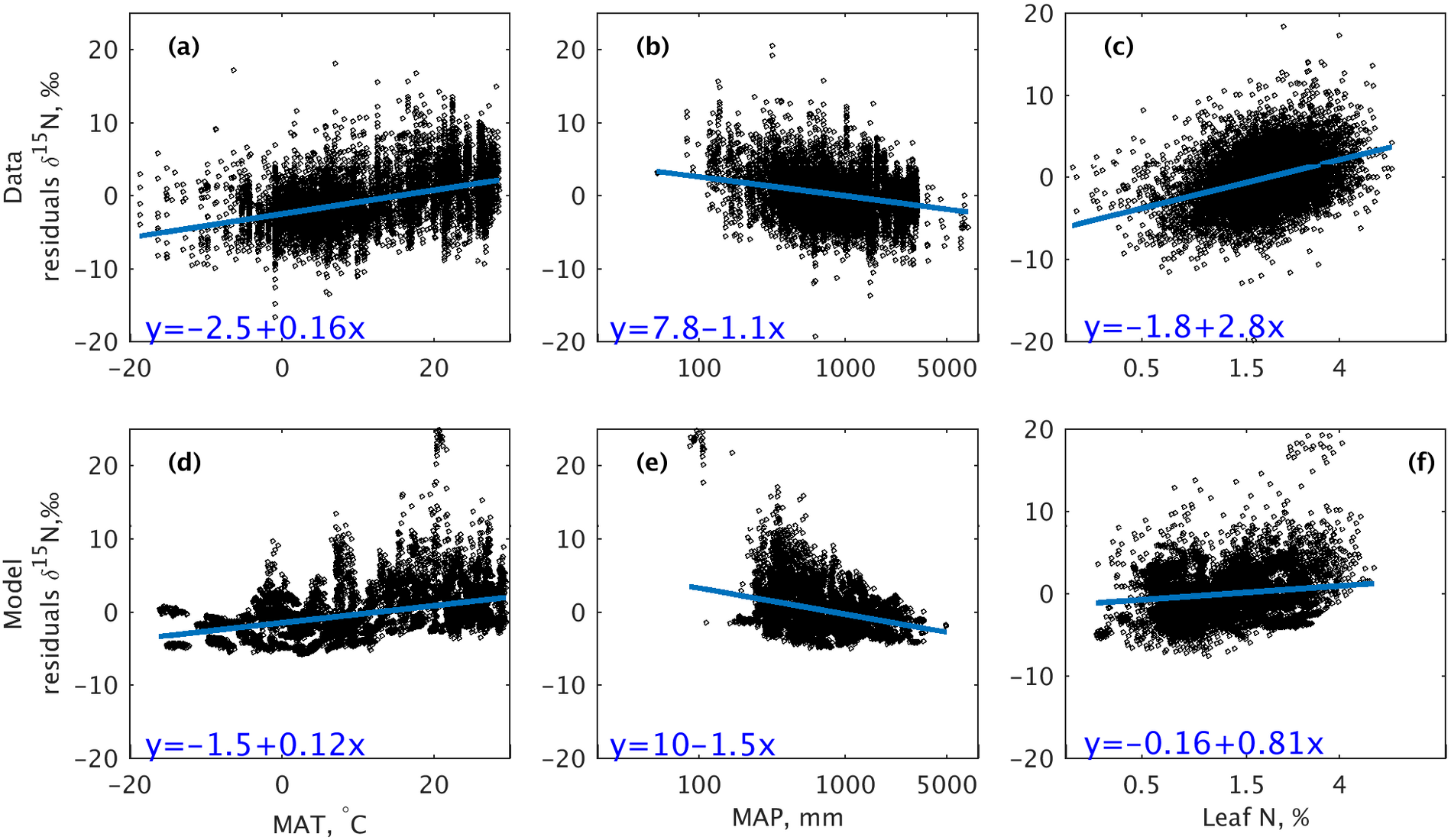
Variation in observed ((a) - (c)) and modelled ((d) - (f)) partial linear regression d15N residuals with mean annual temperature (MAT), mean annual precipitation (MAP) and leaf nitrogen content. Data values are individual observations. Model values are mean leaf d15N residuals for each year. All regression lines shown are significant with p<0.005.

### 3.2 Effect of atmospheric CO_2_ on foliar δ^15^N

Figure 3 shows δ^15^N slopes of regression model residuals in Craine2018 and QUINCY predictions across PFTs. Across the globe, the observed foliar δ^15^N has a stronger negative trend over time than all model runs (−0.041 +/− 0.0^15^ ‰ yr^−1^), with the baseline scenario, driven by changing climate and atmospheric CO_2_, having an average slope close to zero (0.002 +/− 0.0002 ‰ yr^−1^ Fig. 3). The scenario with fixed atmospheric CO_2_ shows on average an increase in δ^15^N (slope 0.0006 +/− 0.00024 ‰ yr^−1^) (Fig. 3(a)). In terms of variation across PFTs, the data show a negative slope across all PFTs, although the slope standard error is relatively large and only the needleleaf seasonal forests show a very strong decrease and a significant slope (−0.22 +/− 3.61e-09 ‰ yr^−1^, p<0.05). In contrast, all predicted model trends have significant slopes (p<0.05) and much lower slope standard errors than the observations.However, only the tropical broadleaf evergreen, needleleaf evergreen and temperate grassland PFTs show significant negative slopes (−0.0019 +/− 0.00013 ‰ yr^−1^ and −0.0013 +/− 0.00014 ‰ yr^−1^ respectively, p<0.05) in the baseline case. The predicted slopes for the fixed CO_2_ scenario are generally more positive than those for the baseline across PFTs, indicating less N limitation in time. The only exception is the needleleaf seasonal forests, where the two slopes have very similar values (0.00081 +/− 0.00017 ‰ yr^−1^ baseline and 0.00085 +/− 0.00017 ‰ yr^−1^ fixed CO_2_, both p<0.05), which would indicate that the change in N limitation predicted by the model for this PFT is driven by other factors than CO_2_.

**Figure 3.**
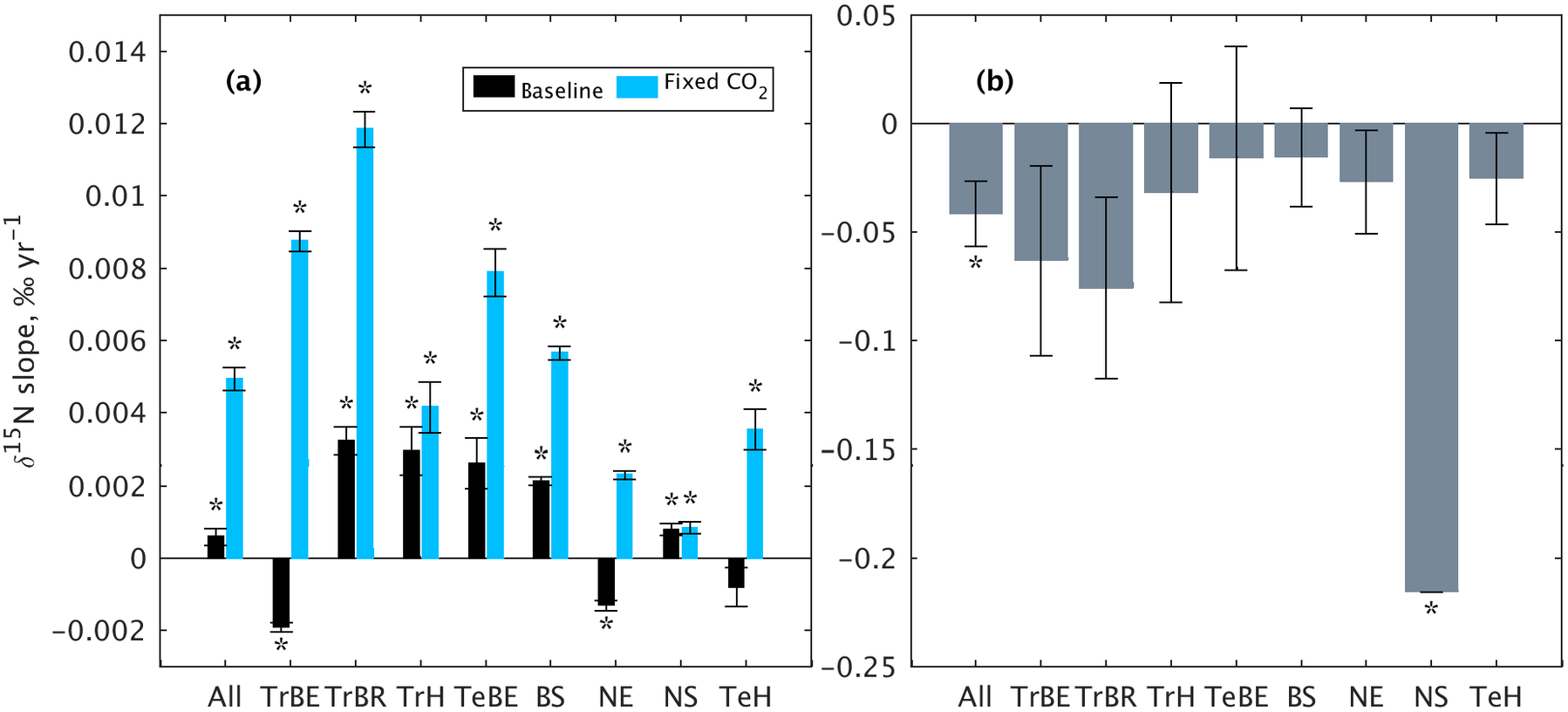
(a) Model d15N residuals slope over time (1980-2018) for the baseline (black) and fixed CO2 (blue) model runs and (b) Craine2018 observed d15N residuals slope for the globe (‘All’, linear regression for all sites) and for each PFT. Error bars represent the standard error of the slope. Stars indicate significant slopes, p<0.05. Model and data shown on different axes for visibility. PFT abbreviations:TrBE tropical broadleaf evergreen,TrBR tropical broadleaf rain green,TrH, C4 grassland, TeBE temperate broadleaf evergreen, BS broadleaf seasonal, NE needleleaf evergreen, NS needleleaf seasonal, TeH C3 grassland.

### 3.3 Effect of atmospheric CO_2_ on ecosystem N status

Figure 4 shows four important ecosystem nutrient status metrics across PFTs for the baseline and fixed CO_2_ model scenarios. In terms of leaf N content (Fig. 4(a)), the baseline model predicts a decrease in all PFTs (mean across all PFTs −0.12 +/− 0.018 % yr^−1^) with the exception of the tropical (C4) grassland, indicating declining nitrogen availability relative to plant growth requirements. Generally, this decrease can be attributed to elevated CO_2_, as it is not predicted by the fixed CO_2_ scenario (mean 0.016 +/− 0.016 % yr^−1^), with the majority of PFTs showing a significant difference between the baseline and the fixed CO_2_ scenarios (p<0.05). However, for needleleaf evergreen forests at higher latitudes, generally considered to be most N limited, there is a slight decrease in simulated leaf N even with no increase in CO_2_, suggesting an increase in N limitation caused by longer growing seasons due to global warming and subsequent increased growth. The tropical grassland is the only PFT where the model predicts an increase in leaf N for both the baseline and the fixed CO_2_ scenarios, with no significant difference between the two, indicating no increase in nitrogen limitation.

**Figure 4.**
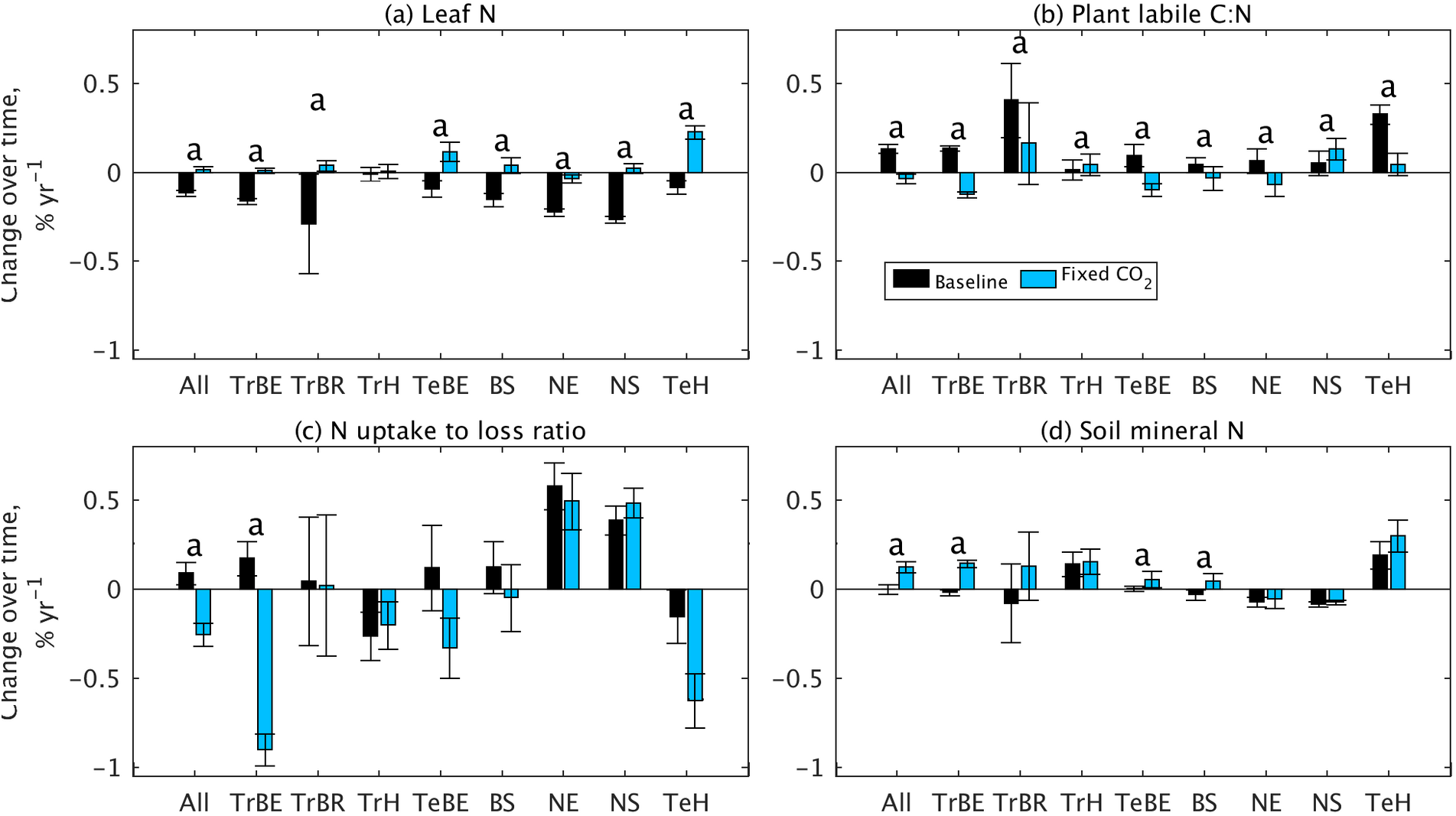
Change in N cycle indicators from 1980 to 2018 across all PFTs due to the increase in atmospheric CO2 as predicted by the model. Relative change in (a) leaf N content, (b) C:N ratio of plant pool available for growth, (c) ratio of plant N uptake to ecosystem loss and (d) soil mineral N. Error bars represent standard error across sites. PFTs for which the model predicts a significant difference between the baseline and fixed CO2 runs are indicated with ‘a’, significance computed with Wilcoxon rank sum test at p<0.05. PFT abbreviations:TrBE tropical broadleaf evergreen,TrBR tropical broadleaf rain-green,TrH, C4 grassland, TeBE temperate broadleaf evergreen, BS broadleaf seasonal, NE needleleaf evergreen, NS needleleaf seasonal, TeHe C3 grassland.

We compare predicted leaf N content with data from the European ICP Forests dataset (Fig. 5), comparing model sites that are located in Europe only. In terms of absolute values, the model predicts slightly lower values than observed on average (observations 1.55 +/− 0.02 %, model 1.27 +/− 0.035 %, Fig. 5(a)), with a stronger bias in the needleleaf evergreen. The model captures the observed change in leaf N content for the broadleaf seasonal (observed −0.23 +/− 0.081 %y^−1^, model −0.18 +/− 0.046 %y^−1^) and needleleaf evergreen (observed −0.29 +/− 0.077 %y^−1^, model −0.22 +/0.02 %y^−1^), while underestimating the change in needleleaf seasonal systems, although the spread in the observed trend in this PFT is relatively large, possibly due to the low number of ICP and model sites in this PFT (Table S3).

**Figure 5.**
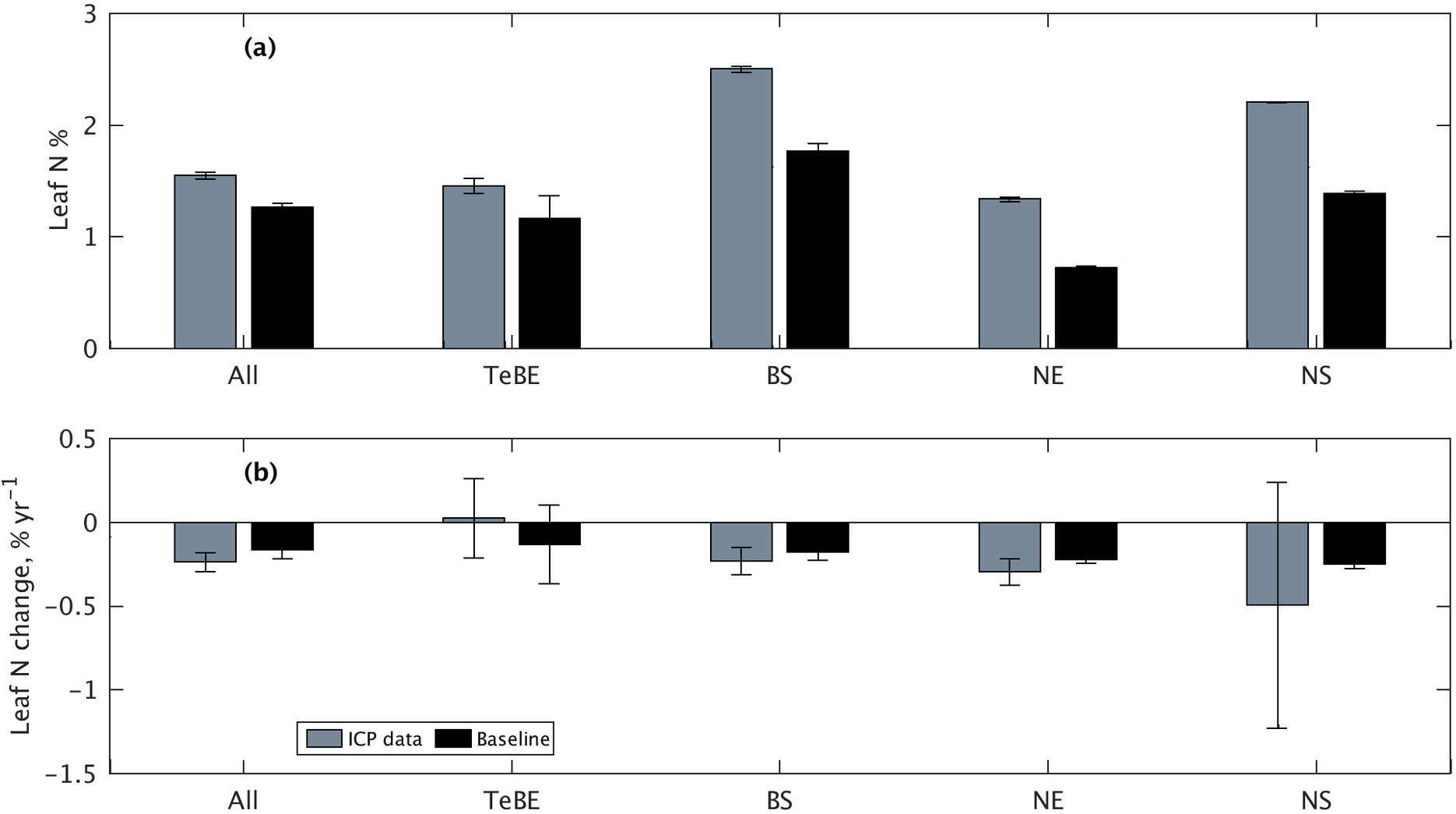
(a) Leaf N content and (b) relative change in leaf N content for 1980 - 2015 observed at the ICP Forests sites and predicted by the model for the baseline model run. Model values are PFT averages in Europe only. Error bars represent standard error across sites.

The C:N ratio of the plant labile pool is another indicator of the nutrient balance of material available for growth and an increase indicates increased nitrogen limitation of plant growth (Fig. 4(b)). The baseline scenario predicts a similar pattern in labile C:N to that seen for leaf N in terms of N limitation, with an increase across all PFTs in the baseline (mean 0.13 +/− 0.024 % yr^−1^) and a small negative change overall for the fixed CO_2_ scenario (−0.037 +/− 0.027 % yr^−1^), with some of the PFTs showing a stronger decrease, implying a decrease in N limitation. There is a significant difference (p<0.05) between the baseline and fixed CO_2_ scenarios in all PFTs, indicating that limitation to growth is driven by increased CO_2_ concentrations. The magnitude of the change between the two plant-based metrics differs across PFTs. For example, the increase in labile C:N is relatively small in the broadleaf seasonal PFT (0.042 +/− 0.04 % yr^−1^), while the response in leaf N magnitude is higher (−0.15 +/− 0.037 % yr^−1^). These patterns indicate differences in plant responses in the different PFTs, so that the model predicts that the change in leaf N content alleviates the limitation to growth. While these are only model predictions, they generally agree with our ecological understanding of the role of flexible leaf stoichiometry. This is important to consider when interpreting leaf N observations as indicators of ecosystem N limitation.

We also examine two measures of whole ecosystem limitation (Fig. 4(c) and (d)). The ratio between plant N uptake to ecosystem N losses is a measure of ‘excess’ N which gets lost from the system when not taken up by plants and an increase in this ratio indicates a tighter N cycle and increased N limitation. Overall, the baseline predicts an increase in this ratio (0.09 +/− 0.063 % yr^−1^), while the fixed CO_2_ scenario predicts a decrease (−0.25 +/− 0.065 % yr^−1^), again indicating that the CO_2_ increase drives ecosystem N cycling further into a N-limited stage. However, the differences between PFTs are very large and the only significant difference between the baseline and fixed CO_2_ runs predicted in the tropical evergreen PFT. The relatively small change in the uptake to loss ratio explains the magnitude in the leaf δ^15^N change (Fig. 3), as also predicted by our conceptual model results (Fig. 1).

In terms of changes in soil mineral N (Fig. 4(d)), on average the baseline model predicts a decrease (−0.0017 +/− 0.027 % yr^−1^) while the fixed CO_2_ scenario predicts an increase (0.13 +/− 0.031 % yr^−1^), although the magnitude of the change is modest in most PFTs. Only the tropical evergreen, temperate evergreen and temperate deciduous PFTs show a significant difference between the two model runs (p<0.05).

### 3.4 Effect of anthropogenic N deposition on foliar δ^15^N trends

We explore our second hypothesis, that changes in the isotopic composition of anthropogenic N deposition is one of the drivers of observed changes in foliar δ^15^N. Overall, both scenarios which include a trend in deposition δ^15^N content (Table 1) predict a lower slope in foliar δ^15^N than the baseline model (Fig. 6), with the strong δ^15^N trend scenario predicting a negative slope (weak δ^15^N trend 0.0011 +/− 0.0011 ‰ yr^−1^, strong δ^15^N trend −0.0064 +/− 0.0011 ‰ yr^−1^), although only the strong δ^15^N trend scenario has a significant slope (p<0.05). Both the overall slopes are less pronounced than the slope in the observations.

**Figure 6.**
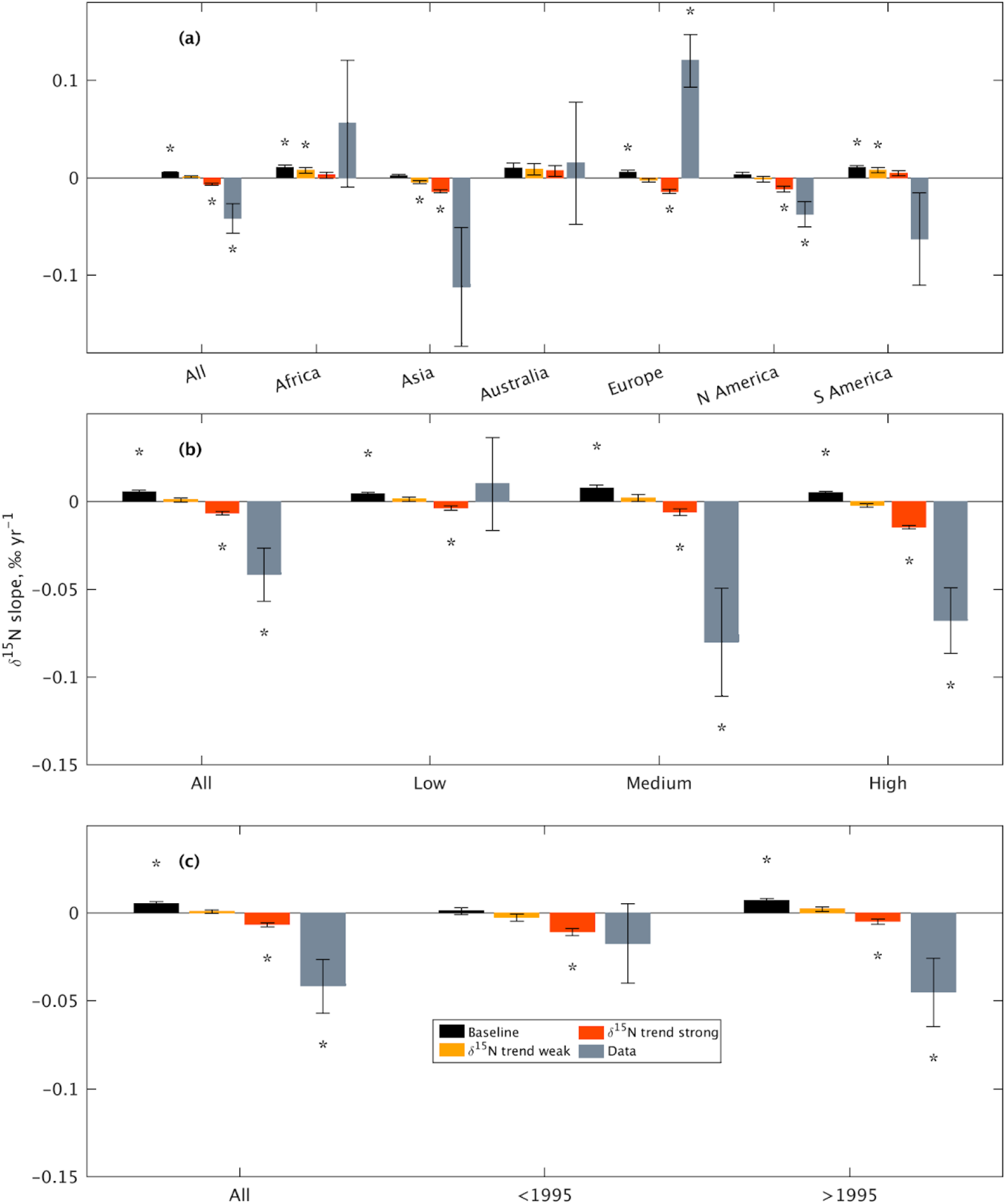
Leaf d15N residuals slope by (a) continent, (b) magnitude of anthropogenic N deposition and (c) timing of peak N deposition. Magnitude classes are: low <0.28 gN m-2yr-1, medium 0.28 - 0.56 gN m-2yr-1 and high >0.56 gN m-2yr-1. Error bars represent slope standard error of the fitted slope. Stars indicate significant slopes, p<0.05

In terms of geographic distribution (Fig. 6(a)) we explore the differences in slope between continents. While the continent separation might seem arbitrary from an ecological point of view, it captures some of the differences in N deposition that come from country-based anthropogenic N emissions levels and subsequent legislation. The observations show a higher variability between continents than that observed between PFTs, with Africa, Australia and Europe having positive slopes (0.056 +/− 0.065 ‰ yr^−1^ and 0.0^15^ +/− 0.063 ‰ yr^−1^ and 0.12 +/− 0.027 ‰ yr^−1^ respectively). However, only Europe and North America show significant trends (p<0.05), reflecting the highly heterogeneous distribution of data availability between continents and in time (Fig. S3). Note here that the majority of observations in Europe are newer than 2000, therefore occurring after the peak of N deposition. While we would expect observations in South America to reflect less N limited tropical and sub-tropical areas, in fact the largest number of measurements are located in the more temperate regions of the continent (Fig. S1). In contrast, our model sites cover the full spectrum of PFTs across the continent and predict a strong positive slope (baseline 0.01 +/− 0.0027 ‰ yr^−1^). As the majority of observations are in the Americas (>60%), the very strong negative slope there has a large effect on the overall slope.

The greatest impact of the simulated δ^15^N trend in deposition occurs in Asia (baseline 0.0021 +/− 0.0018 ‰ yr^−1^, strong δ^15^N trend −0.014 +/− 0.0017 ‰ yr^−1^) and Europe (baseline 0.0059 +/− 0.002 ‰ yr^−1^, strong trend −0.014 +/− 0.002 ‰ yr^−1^). This would on one hand imply an alleviation of N limitation (positive slope) but a greater effect if the N deposition did have a temporal trend in isotopic composition. In contrast, areas with both lower N limitation and N deposition rates (e.g. Africa or Australia) show positive δ^15^N slopes for both the baseline and the model runs with a δ^15^N deposition trend (Fig. 6(a)).

Figure 6 (b) and (c) show the effect of anthropogenic N deposition levels and the timing of peak deposition. The observations show the most negative slopes at sites with medium levels of N deposition (−0.08 +/− 0.031 ‰ yr^−1^) and positive slopes for low deposition sites, although only the medium and high deposition areas have a significant slope (p<0.05). The baseline scenario predicts similar slopes across N deposition ranges. The two scenarios with δ^15^N trend in N deposition predict, as would be expected, a more negative slope with increased N deposition (e.g. low −0.0036 +/− 0.0013 ‰ yr^−1^ to high −0.014 +/− 0.0011 ‰ yr^−1^ for the strong δ^15^N trend run).

In terms of the timing of maximum anthropogenic N deposition (Fig. 6(c)), the observations show a small but not significant slope for sites with an early N deposition peak (−0.016 +/− 0.023 ‰ yr^−1^) and a strongly negative slope (p<0.05) for sites with a more recent peak in N deposition levels (−0.044 +/− 0.019 ‰ yr^−1^). The strong δ^15^N trend model scenario shows a similar pattern with a lower slope for the sites with an early N deposition peak (−0.011 +/− 0.002 ‰ yr^−1^.) compared to the later peaking sites (−0.0047 +/− 0.0014 ‰ yr^−1^). Both of these groups have significant slopes in the strong δ^15^N trend scenario while for the baseline and weak trend scenarios, only the late peaking sites show significant slopes (p<0.05).

## 4 Discussion

We compared predictions from an isotope enabled LSM with a global, long-term dataset of foliar δ^15^N. We demonstrated that the model captures large-scale trends in foliar dN^15^ associated with mean annual temperature, precipitation and foliar N recorded in the data. The model furthermore reproduces trends in foliar N concentrations across European forest ecosystems, for which we had sufficient data available to assess the simulated trends. However, the model does not simulate the strong decreasing trend in foliar δ^15^N present in the observations. Consistent with hypothesis 1, QUINCY shows an overall lower leaf δ^15^N with elevated CO_2_ compared to the model scenario with fixed CO_2_, even if the magnitude of the slope is much smaller than the observation-derived trend. Using idealised scenarios, we demonstrate isotopic changes in N deposition, a phenomenon unrelated to nutrient limitation, may provide a stronger contribution to the decreased trend in foliar δ^15^N than changes in soil N availability. Our results point to difficulty of identifying the causal processes between observed variations in foliar δ^15^N, especially at large spatial and temporal scales and therefore caution against interpretation of the observed trend in foliar ^15^N as indicative of large-scale oligotrophication due to rising atmospheric CO_2_.

### 4.1 Nitrogen limitation with elevated CO_2_

We hypothesised that increases in atmospheric CO_2_ lead to increased ecosystem N limitation. Indeed, the model predicts N limitation across most PFTs (Fig. 4) and captures observed decreases in leaf N only for the model run with transient CO_2_. Increasing N limitation, or rather declining N availability, with increasing CO_2_ is associated with a stronger decline in foliar δ^15^N than when keeping CO_2_ at pre-industrial levels. However, the magnitude of this trend is substantially smaller than the trend inferred from observations. Overall, our results confirm the hypothesis, but the process interaction is complex: while the decrease in leaf N occurs under transient CO_2_ only, this does not always translate into enhanced growth limitation (Fig. 4(b) and S5). This can be explained by two factors: (1) decrease in N content is a photosynthetic response at the leaf level (Ainsworth & Long, 2005) and (2) decrease in N content alleviates limitation and allows plants to maintain growth. Therefore, a decrease in leaf N or δ^15^N is a sign of reduced N availability but not necessarily increased growth limitation. The exact relationship between soil N availability (Fig. 4(c) and (d)), limitation to growth (Fig. 4(b) and S5) and changes in leaf N and δ^15^N (Fig. 4(a) and 3(a)) varies across PFTs and implicitly with plant strategies and adaptations in real world observations.

### 4.2 Model and data uncertainties

To our knowledge, QUINCY is the first LSM to include nitrogen isotope fractionation. This can be a powerful tool for evaluating N cycle processes, both by using N tracer experiments and natural abundance data. Here, QUINCY broadly reproduces expected patterns of foliar δ^15^N, both at process level (Fig.1) and at broader spatial scales (Fig. 2). This supports the application of the model to analyse trends in foliar ^15^N and compare these to observations.

We do however observe a mismatch between observed and predicted temporal trends in leaf δ^15^N, in that data shows a much stronger decrease than the model (Figs. 3 and 6). There can be multiple causes behind this mismatch, if we assume isotope representation in QUINCY is correct: (1) other processes acting alongside N limitation, (2) model representation of N cycle processes and (3) a difference in sites covered by the observations and by the model respectively. The first point refers to our second hypothesis that changes in anthropogenic N deposition affect the foliar δ^15^N signal. Our results support this hypothesis, in that model simulations which include a δ^15^N trend in N deposition predict a negative slope in δ^15^N on average, with model predictions matching observations more closely in areas with higher N deposition, although there is still discrepancy between observed and modelled trends (Fig. 6). Another supporting argument is that observed trends are connected to timing of N deposition and therefore changes in legislation and deposition sources. Areas with relatively high levels of N deposition (Asia, North America) show a strong observed negative trend ( Fig 6(a)), although we would expect an alleviation of N limitation by the N input. Europe, the area with the highest historical levels of N deposition, shows a significant positive trend in the observations, although this might be explained by the skewed temporal distribution of observations here, with most available data being after 2000 (Fig. S3), while the peak in N deposition in most European countries was in the 1980s (Engardt et al., 2017). Additionally, as different sources of deposition have different isotopic signatures (Elliott et al., 2019), the exact trend can depend on the spatial and temporal distribution of emission sources.

Overall, the evidence for our second hypothesis is mixed. The model scenario with a strong trend in δ^15^N in N deposition is the only one to predict a negative global slope of foliar δ^15^N, but the magnitude is still lower than the observations. As discussed above, the continentally segregated spatial patterns found in the observations somewhat support the hypothesis but multiple factors affecting δ^15^N signals make attribution difficult. Our implementation of the deposition trend was based on two studies (Hastings et al., 2009; Holtgrieve et al., 2011) which aim to represent large-scale temporal patterns in δ^15^N. We chose to implement values from both studies as the difference in observed changes is relatively large and we aimed to represent different possible scenarios. While we would generally expect δ^15^N signatures of deposition to be less spatially and temporally heterogeneous than total N deposited (Bauer et al., 2000), there is still some variability. Both studies are based in the northern hemisphere and while atmospheric transport should mean lake sediments and ice cores can represent signals from large areas of the globe, it is unrealistic that observed trends are fully representative of a global process. Differences between the two records could be caused by fractionation processes e.g. in lake sediments (Lehmann et al., 2003; Leng et al., 2006). beyond the scope of this paper. Further, such processes could dilute the deposition δ^15^N signal, so the trend we use here might be weaker than what would actually be observed in deposition. It is equally unrealistic that a single trend can be applied to a set of globally distributed sites, an approach which we have taken due to a lack of data. However, we show that a temporal trend in the ^15^N content of N deposition does have an effect on foliar δ^15^N and should therefore be taken into consideration when looking at long term trends. To correctly match the spatial distribution of observations, we would need a more spatially distributed dataset of δ^15^N content of N deposition and its change over time.

An alternative explanation of the mismatch between model predictions and observations is that QUINCY does not correctly predict an increase in N limitation. However, the simulations show an increase in N limitation based on all other metrics used in this study (Fig. 4) and the magnitude of the response is correct compared to observed changes in leaf N content across European forests (Fig. 5). A more comprehensive, global dataset of leaf N timeseries would allow a more extensive global evaluation, but even with the limited data available, QUINCY predicts decreases in leaf N over time within observed ranges.

Another possible reason behind the data-model mismatch is the sites covered by the model and the observations. We designed our study to represent sites distributed across climate zones and PFTs to represent a global picture, rather than the site distribution present in the Craine2018 dataset (Fig. S1 and S2). Additionally, the temporal distribution of observations is highly skewed towards the present day. This results in a leaf δ^15^N trend representative of the higher latitudes and the last 20 years. Additionally, when we partition the data into smaller regions, even though resulting δ^15^N slopes are more negative than those predicted by the model, the trends are very heterogeneous and very few slopes are actually statistically significant, due to the multiple processes contributing to N isotope fractionation. Craine2018 is the best dataset of foliar δ^15^N we have to date and a coordinated observation network is needed to address these gaps.

### 4.3 Use of foliar δ^15^N data for assessing ecosystem N limitation status

Natural abundance δ^15^N data can be a very powerful tool in investigating ecosystem N cycle processes. However, the main advantage that plant δ^15^N integrates many processes and can therefore be representative of ecosystem level N status, can also be seen as its main disadvantage, when many processes contribute to small changes it becomes difficult to attribute causality (Craine et al., 2015). For natural abundance studies such as ours, the relationship between N availability and foliar δ^15^N at site-level can be explained in a relatively straightforward manner (Fig. 1). Once we add the dimensions of time and space, the problem becomes increasingly complex. Although we remove average effects of climate (temperature and precipitation), there are still a large number of variables controlling isotope fractionation processes, including N availability before anthropogenic disturbance, plant functional type and N deposition amount and temporal patterns. An ideal dataset for assessing changes in ecosystem N limitation in time would be a consistent timeseries of observations at the same sites, including measurements of soil and leaf δ^15^N following a common protocol, similar to the kind of information available from the ICP Forests database. In addition to leaf N isotope content, δ^15^N in tree rings can provide a consistent timeseries of N availability (Martin et al., 2021; van der Sleen et al., 2015) and a distributed database of such measurements could prove to be an extremely powerful tool.

Independent of how robustly we could identify a decrease in soil N availability over time from a δ^15^N dataset, the question of plant growth limitation still remains. Our model predicts a decrease in leaf N with increased CO_2_ across almost all PFTs (Fig. 4(a)) but the change in N available to growth is neither as prevalent nor as strong as that in leaf N (Fig. 4(b) and S5). This would imply that plants adjust their stoichiometry under decreased N availability to maximise productivity thus alleviating N limitation, a hypothesis that has been advanced before (Drake et al., 1997). While in the current study we cannot evaluate this model prediction, it does indicate that foliar δ^15^N data should be used alongside plant biomass data to actually infer the effect of changes in soil N on vegetation growth.

### 4.4 Combining models with data to understand patterns of N limitation

One of our aims in this study was to showcase the usefulness of an LSM alongside a large observational dataset to interpret observed patterns and for process attribution. An LSM allows us to simulate different scenarios and to ‘turn off’ processes to test hypotheses. We show that in the absence of any CO_2_ increase QUINCYpredicts a higher foliar δ^15^N, although the pattern differs among PFTs. For example, tropical regions show a lower N limitation increase with increasing CO_2_, likely due to high levels of N fixation in these systems (Cleveland et al., 1999). On the other hand, boreal regions show a decrease in δ^15^N even without an increase in CO_2_. While geographical patterns could be inferred simply from data, the effect of CO_2_ could not. Another advantage of using an LSM is that we can derive quantities that are not commonly measured, such as internal plant N availability and soil N cycling, to further investigate ecosystem N limitation and draw conclusions about vegetation N status.

Our analysis highlights two important points: (1) the importance of N deposition for ecosystem N limitation, not only as N inputs but also through the way its isotopic composition affects our interpretation of plant and soil δ^15^N and (2) the difference in N limitation viewed through different ecosystem metrics (Fig. 4). The first point was discussed in section 4.1, but in summary the ^15^N signal of anthropogenic N deposition input to ecosystems affects foliar δ^15^N, so that the temporal and spatial pattern of N emission sources must be taken into account when analysing long-term foliar δ^15^N data. While a readily available data product does not yet exist, some of the pieces needed to start building such a product do (emission inventories, atmospheric transport models).

The complexity of N limitation metrics is an issue acknowledged before (Vicca et al., 2018), as we ourselves describe in the introduction, but the model analysis paints a clearer picture of what the processes are. A decrease in leaf N, as predicted by QUINCY in response to increased CO_2_ is not always a good indicator of growth limitation, as a decrease either alleviates the limitation or is a photosynthesis level response to elevated CO_2_. Soil mineral N is also not a very good indicator, due to the many plant and microbial processes that interact with it. While CN stoichiometry of the plant labile pool is the most direct metric of how much plant growth is actually N limited, this is unfortunately very difficult to measure, if at all possible. However, if data were available at a network of sites, for both leaf and soil N content, as well as soil N loss, these measures could be used to evaluate the model after which predicted labile CN could be used with confidence to determine the N limitation status.

Overall, we conclude that large datasets of leaf nitrogen and its isotopic composition are not, on their own, sufficient to assess changes in ecosystem N status. We show temporal trends in leaf δ^15^N are not exclusively driven by oligotrophication and a decrease in leaf N content does not always translate intoN limitation to plant growth. Foliar concentrations result from a complex chain of soil and plant processes and are therefore difficult to compare across sites without additional information. e propose to go beyond using data to evaluate models and to use models and data jointly to further interrogate ecosystem processes and draw conclusions about possible, difficult to observe mechanisms, as well as identify data gaps and future paths to further our understanding of the processes at hand.

## Acknowledgments

This work was supported by the European Research Council (ERC) under the European Union’s Horizon 2020 research and innovation programme (QUINCY; grant no. 647204). LY was supported by the Swedish government funded Strategic Research Area Biodiversity and Ecosystems in a Changing Climate, BECC. TT was supported by the Academy of Finland (grant no. 330165). The evaluation was based on data that was collected by partners of the official UNECE ICP Forests Network (http://icp-forests.net/contributors). Part of the data was co-financed by the European Commission. We are grateful to Dr. Jan Engel for technical assistance in developing the code.

## Author contributions

SC and SZ designed the study. SC performed the analyses. TT contributed to initial exploratory analysis. RN contributed to statistical analyses and result interpretation. SC, TT, LY, MK and SZ developed the model that forms the basis of the analysis in this manuscript. All authors contributed to writing the manuscript.

## Data availability

The global dataset of leaf δ^15^N observation is freely available at https://doi.org/10.5061/dryad.v2k2607. The ICP Forests database is available by request from http://icp-forests.net/.

## Code availability

The scientific part of the code is available under a GPL v3 licence. The scientific code of QUINCY relies on software infrastructure from the MPI-ESM environment, which is subject to the MPI-M-Software-License-Agreement in its most recent form (http://www.mpimet.mpg.de/en/science/models/license). The source code is available online (https://git.bgc-jena.mpg.de/quincy/quincy-model-releases), but its access is restricted to registered users. Readers interested in running the model should request a username and password from the corresponding authors or via the git-repository. Model users are strongly encouraged to follow the fair-use policy stated on https://www.bgc-jena.mpg.de/bgi/index.php/Projects/QUINCYModel.

## References

Ainsworth, E. A., & Long, S. P. (2005). What have we learned from 15 years of free-air CO2 enrichment (FACE)? A meta-analytic review of the responses of photosynthesis, canopy properties and plant production to rising CO2. The New Phytologist, 165(2), 351–371. https://doi.org/10.1111/j.1469-8137.2004.01224.x

Amundson, R., Austin, A. T., Schuur, E. A. G., Yoo, K., Matzek, V., Kendall, C., Uebersax, A., Brenner, D., & Baisden, W. T. (2003). Global patterns of the isotopic composition of soil and plant nitrogen. Global Biogeochemical Cycles, 17(1). https://agupubs.onlinelibrary.wiley.com/doi/abs/10.1029/2002gb001903

Arora, V. K., Katavouta, A., Williams, R. G., Jones, C. D., Brovkin, V., Friedlingstein, P., Schwinger, J., Bopp, L., Boucher, O., Cadule, P., Chamberlain, M. A., Christian, J. R., Delire, C., Fisher, R. A., Hajima, T., Ilyina, T., Joetzjer, E., Kawamiya, M., Koven, C. D., … Ziehn, T. (2020). Carbon–concentration and carbon–climate feedbacks in CMIP6 models and their comparison to CMIP5 models. Biogeosciences, 17(16), 4173–4222. https://doi.org/10.5194/bg-17-4173-2020

Atkin, O. K., Bloomfield, K. J., Reich, P. B., Tjoelker, M. G., Asner, G. P., Bonal, D., Bönisch, G., Bradford, M. G., Cernusak, L. A., Cosio, E. G., Creek, D., Crous, K. Y., Domingues, T. F., Dukes, J. S., Egerton, J. J. G., Evans, J. R., Farquhar, G. D., Fyllas, N. M., Gauthier, P. P. G., … Zaragoza-Castells, J. (2015). Global variability in leaf respiration in relation to climate, plant functional types and leaf traits. The New Phytologist, 206(2), 614–636. https://doi.org/10.1111/nph.13253

Bauer, G. A., Gebauer, G., Harrison, A. F., Högberg, P., Högbom, L., Schinkel, H., Taylor, A. F. S., Novak, M., Buzek, F., Harkness, D., Persson, T., & Schulze, E.-D. (2000). Biotic and Abiotic Controls Over Ecosystem Cycling of Stable Natural Nitrogen, Carbon and Sulphur Isotopes. In E.-D. Schulze (Ed.), Carbon and Nitrogen Cycling in European Forest Ecosystems(pp. 189–214). Springer Berlin Heidelberg. https://doi.org/10.1007/978-3-642-57219-7_9

Caldararu, S., Thum, T., Yu, L., & Zaehle, S. (2020). Whole-plant optimality predicts changes in leaf nitrogen under variable CO 2 and nutrient availability. The New Phytologist, 225(6), 2331–2346. https://nph.onlinelibrary.wiley.com/doi/abs/10.1111/nph.16327

Choi, W.-J., Chang, S. X., Allen, H. L., Kelting, D. L., & Ro, H.-M. (2005). Irrigation and fertilization effects on foliar and soil carbon and nitrogen isotope ratios in a loblolly pine stand. Forest Ecology and Management, 213(1), 90–101. https://doi.org/10.1016/j.foreco.2005.03.016

Cleveland, C. C., Townsend, A. R., Schimel, D. S., Fisher, H., Howarth, R. W., Hedin, L. O., Perakis, S. S., Latty, E. F., Von Fischer, J. C., Elseroad, A., & Others. (1999). Global patterns of terrestrial biological nitrogen (N2) fixation in natural ecosystems. Global Biogeochemical Cycles, 13(2), 623–645. https://onlinelibrary.wiley.com/doi/pdf/10.1029/1999GB900014

Craine, J. M., Brookshire, E. N. J., Cramer, M. D., Hasselquist, N. J., Koba, K., Marin-Spiotta, E., & Wang, L. (2015). Ecological interpretations of nitrogen isotope ratios of terrestrial plants and soils. Plant and Soil, 396(1), 1–26. https://doi.org/10.1007/s11104-015-2542-1

Craine, J. M., Elmore, A. J., Aidar, M. P. M., Bustamante, M., Dawson, T. E., Hobbie, E. A., Kahmen, A., Mack, M. C., McLauchlan, K. K., Michelsen, A., Nardoto, G. B., Pardo, L. H., Peñuelas, J., Reich, P. B., Schuur, E. A. G., Stock, W. D., Templer, P. H., Virginia, R. A., Welker, J. M., & Wright, I. J. (2009). Global patterns of foliar nitrogen isotopes and their relationships with climate, mycorrhizal fungi, foliar nutrient concentrations, and nitrogen availability. The New Phytologist, 183(4), 980–992. https://doi.org/10.1111/j.1469-8137.2009.02917.x

Craine, J. M., Elmore, A. J., Wang, L., Aranibar, J., Bauters, M., Boeckx, P., Crowley, B. E., Dawes, M. A., Delzon, S., Fajardo, A., Fang, Y., Fujiyoshi, L., Gray, A., Guerrieri, R., Gundale, M. J., Hawke, D. J., Hietz, P., Jonard, M., Kearsley, E., … Zmudczyńska-Skarbek, K. (2018). Isotopic evidence for oligotrophication of terrestrial ecosystems. Nature Ecology & Evolution, 2(11), 1735–1744. https://doi.org/10.1038/s41559-018-0694-0

Davies-Barnard, T., Meyerholt, J., Zaehle, S., Friedlingstein, P., Brovkin, V., Fan, Y., Fisher, R. A., Jones, C. D., Lee, H., Peano, D., Smith, B., Wårlind, D., & Wiltshire, A. J. (2020).Nitrogen cycling in CMIP6 land surface models: progress and limitations [Data set]. https://doi.org/10.5194/bg-17-5129-2020

Drake, B. G., Gonzalez-Meler, M. A., & Long, S. P. (1997). More Efficient Plants: A Consequence of Rising Atmospheric CO2? Annual Review of Plant Physiology and Plant Molecular Biology, 48, 609–639. https://doi.org/10.1146/annurev.arplant.48.1.609

Elliott, E. M., Yu, Z., Cole, A. S., & Coughlin, J. G. (2019). Isotopic advances in understanding reactive nitrogen deposition and atmospheric processing. The Science of the Total Environment, 662, 393–403. https://doi.org/10.1016/j.scitotenv.2018.12.177

Ellsworth, D. S., Reich, P. B., Naumburg, E. S., Koch, G. W., Kubiske, M. E., & Smith, S. D. (2004). Photosynthesis, carboxylation and leaf nitrogen responses of 16 species to elevated pCO2 across four free-air CO2 enrichment experiments in forest, grassland and desert. Global Change Biology, 10(12), 2121–2138. https://doi.org/10.1111/j.1365-2486.2004.00867.x

Elser, J. J., Fagan, W. F., Kerkhoff, A. J., Swenson, N. G., & Enquist, B. J. (2010). Biological stoichiometry of plant production: metabolism, scaling and ecological response to global change. The New Phytologist, 186(3), 593–608. https://doi.org/10.1111/j.1469-8137.2010.03214.x

Engardt, M., Simpson, D., Schwikowski, M., & Granat, L. (2017). Deposition of sulphur and nitrogen in Europe 1900–2050. Model calculations and comparison to historical observations. Tellus. Series B, Chemical and Physical Meteorology, 69(1), 1328945. https://doi.org/10.1080/16000889.2017.1328945

Felix, J. D., David Felix, J., Elliott, E. M., & Shaw, S. L. (2012). Nitrogen Isotopic Composition of Coal-Fired Power Plant NOx: Influence of Emission Controls and Implications for Global Emission Inventories. In Environmental Science & Technology (Vol. 46, Issue 6, pp. 3528–3535). https://doi.org/10.1021/es203355v

Fowler, D., Coyle, M., Skiba, U., Sutton, M. A., Cape, J. N., Reis, S., Sheppard, L. J., Jenkins, A., Grizzetti, B., Galloway, J. N., Vitousek, P., Leach, A., Bouwman, A. F., Butterbach-Bahl, K., Dentener, F., Stevenson, D., Amann, M., & Voss, M. (2013). The global nitrogen cycle in the twenty-first century. Philosophical Transactions of the Royal Society of London. Series B, Biological Sciences, 368(1621), 20130164. https://doi.org/10.1098/rstb.2013.0164

Friend, A. D. (2010). Terrestrial plant production and climate change. Journal of Experimental Botany, 61(5), 1293–1309. https://doi.org/10.1093/jxb/erq019

Galloway, J. N., Dentener, F. J., Capone, D. G., Boyer, E. W., Howarth, R. W., Seitzinger, S. P., Asner, G. P., Cleveland, C. C., Green, P. A., Holland, E. A., & Others. (2004). Nitrogen cycles: past, present, and future. Biogeochemistry, 70(2), 153–226. https://idp.springer.com/authorize/casa?redirect_uri=https://link.springer.com/article/10.1007/s10533-004-0370-0&casa_token=hZ0aXggRZ54AAAAA:Z5oUS-Tjxgc0-n0BLLwoo_8j7gNXdXv9a5BvCu9QxOlXiasCW2u_Yuo0gERIVyrwdQjL7754Km0-voOH

Garten, C. T., Jr, & Van Miegroet, H. (1994). Relationships between soil nitrogen dynamics and natural ^15^N abundance in plant foliage from Great Smoky Mountains National Park. Canadian Journal of Forest Research. Journal Canadien de La Recherche Forestiere, 24(8), 1636–1645. https://www.nrcresearchpress.com/doi/abs/10.1139/x94-212

Harris, I. C. (2020). CRU JRA v2.1: A forcings dataset of gridded land surface blend of Climatic Research Unit (CRU) and Japanese reanalysis (JRA) data; Jan.1901 - Dec.2019.. Centre for Environmental Data Analysis[Data set].

Hastings, M. G., Jarvis, J. C., & Steig, E. J. (2009). Anthropogenic impacts on nitrogen isotopes of ice-core nitrate. Science, 324(5932), 1288. https://doi.org/10.1126/science.1170510

Hengl, T., Mendes de Jesus, J., Heuvelink, G. B. M., Ruiperez Gonzalez, M., Kilibarda, M., Blagotić, A., Shangguan, W., Wright, M. N., Geng, X., Bauer-Marschallinger, B., Guevara, M. A., Vargas, R., MacMillan, R. A., Batjes, N. H., Leenaars, J. G. B., Ribeiro, E., Wheeler, I., Mantel, S., & Kempen, B. (2017). SoilGrids250m: Global gridded soil information based on machine learning. PloS One, 12(2), e0169748. https://doi.org/10.1371/journal.pone.0169748

Hogberg, P. (1998). Tansley Review No. 95: ^15^N natural abundance in soil-plant systems. In New Phytologist (Vol. 139, Issue 3, pp. 595–595). https://doi.org/10.1046/j.1469-8137.1998.00239.x

Holtgrieve, G. W., Schindler, D. E., Hobbs, W. O., Leavitt, P. R., Ward, E. J., Bunting, L., Chen, G., Finney, B. P., Gregory-Eaves, I., Holmgren, S., Lisac, M. J., Lisi, P. J., Nydick, K., Rogers, L. A., Saros, J. E., Selbie, D. T., Shapley, M. D., Walsh, P. B., & Wolfe, A. P. (2011). A coherent signature of anthropogenic nitrogen deposition to remote watersheds of the Northern Hemisphere. Science, 334(6062), 1545–1548. https://doi.org/10.1126/science.1212267

Hungate, B. A., Dukes, J. S., Shaw, M. R., Luo, Y., & Field, C. B. (2003). Atmospheric science. Nitrogen and climate change. Science, 302(5650), 1512–1513. https://doi.org/10.1126/science.1091390

Hurtt, G. C., Frolking, S., Fearon, M. G., Moore, B., Shevliakova, E., Malyshev, S., Pacala, S. W., & Houghton, R. A. (2006). The underpinnings of land-use history: three centuries of global gridded land-use transitions, wood-harvest activity, and resulting secondary lands. Global Change Biology, 12(7), 1208–1229. https://doi.org/10.1111/j.1365-2486.2006.01150.x

Johannisson, C., & Högberg, P. (1994). ^15^N abundance of soils and plants along an experimentally induced forest nitrogen supply gradient. Oecologia, 97(3), 322–325. https://doi.org/10.1007/BF00317321

Jonard, M., Fürst, A., Verstraeten, A., Thimonier, A., Timmermann, V., Potočić, N., Waldner, P., Benham, S., Hansen, K., Merilä, P., Ponette, Q., de la Cruz, A. C., Roskams, P., Nicolas, M., Croisé, L., Ingerslev, M., Matteucci, G., Decinti, B., Bascietto, M., & Rautio, P. (2015). Tree mineral nutrition is deteriorating in Europe. Global Change Biology, 21(1), 418–430. https://doi.org/10.1111/gcb.12657

Jones, C. D., & Friedlingstein, P. (2020). Quantifying process-level uncertainty contributions to TCRE and carbon budgets for meeting Paris Agreement climate targets. Environmental Research Letters: ERL [Web Site], 15(7), 074019. https://doi.org/10.1088/1748-9326/ab858a

Kattge, J., Bönisch, G., Díaz, S., Lavorel, S., Prentice, I. C., Leadley, P., Tautenhahn, S., Werner, G. D. A., Aakala, T., Abedi, M., Acosta, A. T. R., Adamidis, G. C., Adamson, K., Aiba, M., Albert, C. H., Alcántara, J. M., Alcázar C, C., Aleixo, I., Ali, H., … Wirth, C. (2020). TRY plant trait database - enhanced coverage and open access. Global Change Biology, 26(1), 119–188. https://doi.org/10.1111/gcb.14904

Kull, O., & Kruijt, B. (1998). Leaf photosynthetic light response: a mechanistic model for scaling photosynthesis to leaves and canopies. Functional Ecology, 12(5), 767–777. https://doi.org/10.1046/j.1365-2435.1998.00257.x

Lamarque, J.-F., Bond, T. C., Eyring, V., Granier, C., Heil, A., Klimont, Z., Lee, D., Liousse, C., Mieville, A., Owen, B., Schultz, M. G., Shindell, D., Smith, S. J., Stehfest, E., Van Aardenne, J., Cooper, O. R., Kainuma, M., Mahowald, N., McConnell, J. R., … van Vuuren, D. P. (2010). Historical (1850–2000) gridded anthropogenic and biomass burning emissions of reactive gases and aerosols: methodology and application. Atmospheric Chemistry and Physics, 10(15), 7017–7039. https://doi.org/10.5194/acp-10-7017-2010

Lamarque, J.-F., Kyle, G. P., Meinshausen, M., Riahi, K., Smith, S. J., van Vuuren, D. P., Conley, A. J., & Vitt, F. (2011). Global and regional evolution of short-lived radiatively-active gases and aerosols in the Representative Concentration Pathways. Climatic Change, 109(1), 191. https://doi.org/10.1007/s10584-011-0155-0

Lehmann, M. F., Reichert, P., Bernasconi, S. M., Barbieri, A., & McKenzie, J. A. (2003). Modelling nitrogen and oxygen isotope fractionation during denitrification in a lacustrine redox-transition zone. Geochimica et Cosmochimica Acta, 67(14), 2529–2542. https://doi.org/10.1016/S0016-7037(03)00085-1

Leng, M. J., Lamb, A. L., Heaton, T. H. E., Marshall, J. D., Wolfe, B. B., Jones, M. D., Holmes, J. A., & Arrowsmith, C. (2006). ISOTOPES IN LAKE SEDIMENTS. In M. J. Leng (Ed.), Isotopes in Palaeoenvironmental Research (pp. 147–184). Springer Netherlands. https://doi.org/10.1007/1-4020-2504-1_04

Le Quéré, C., Andrew, R. M., Friedlingstein, P., Sitch, S., Hauck, J., Pongratz, J., Pickers, P. A., Korsbakken, J. I., Peters, G. P., Canadell, J. G., Arneth, A., Arora, V. K., Barbero, L., Bastos, A., Bopp, L., Chevallier, F., Chini, L. P., Ciais, P., Doney, S. C., … Zheng, B. (2018). Global Carbon Budget 2018. Earth System Science Data, 10(4), 2141–2194. https://doi.org/10.5194/essd-10-2141-2018

Lorenz, M. (1995). International Co-operative Programme on Assessment and Monitoring of Air Pollution Effects on Forests-ICP Forests-. Water, Air, and Soil Pollution, 85(3), 1221–1226. https://doi.org/10.1007/BF00477148

Magill, A. H., Aber, J. D., Currie, W. S., Nadelhoffer, K. J., Martin, M. E., McDowell, W. H., Melillo, J. M., & Steudler, P. (2004). Ecosystem response to 15 years of chronic nitrogen additions at the Harvard Forest LTER, Massachusetts, USA. Forest Ecology and Management, 196(1), 7–28. https://doi.org/10.1016/j.foreco.2004.03.033

Martin, A. C., Macias-Fauria, M., Bonsall, M. B., Forbes, B. C., Zetterberg, P., & Jeffers, E. S. (2021). Common mechanisms explain nitrogen-dependent growth of Arctic shrubs over three decades despite heterogeneous trends and declines in soil nitrogen availability. The New Phytologist, nph.17529. https://doi.org/10.1111/nph.17529

Matson, P., Lohse, K. A., & Hall, S. J. (2002). The globalization of nitrogen deposition: consequences for terrestrial ecosystems. Ambio, 31(2), 113–119. https://doi.org/10.1579/0044-7447-31.2.113

Meyerholt, J., Zaehle, S., & Smith, M. J. (2016). Variability of projected terrestrial biosphere responses to elevated levels of atmospheric CO2 due to uncertainty in biological nitrogen fixation. Biogeosciences, 13(5), 1491–1518. https://doi.org/10.5194/bg-13-1491-2016

Parton, W. J., Scurlock, J. M. O., Ojima, D. S., Gilmanov, T. G., Scholes, R. J., Schimel, D. S., Kirchner, T., Menaut, J.-C., Seastedt, T., Garcia Moya, E., Kamnalrut, A., & Kinyamario, J. I. (1993). Observations and modeling of biomass and soil organic matter dynamics for the grassland biome worldwide. Global Biogeochemical Cycles, 7(4), 785–809. https://doi.org/10.1029/93GB02042

Rastetter, E. B., Vitousek, P. M., Field, C., Shaver, G. R., Herbert, D., & Gren, G. I. (2001). Resource Optimization and Symbiotic Nitrogen Fixation. Ecosystems, 4(4), 369–388. https://doi.org/10.1007/s10021-001-0018-z

Robinson, D. (2001). δ^15^N as an integrator of the nitrogen cycle. Trends in Ecology & Evolution, 16(3), 153–162. https://doi.org/10.1016/S0169-5347(00)02098-X

Sikström, U. (2002). Effects of liming and fertilization (N, PK) on stem growth, crown transparency, and needle element concentrations of Picea abies stands in southwestern Sweden. Canadian Journal of Forest Research. Journal Canadien de La Recherche Forestiere, 32(10), 1717–1727. https://doi.org/10.1139/x02-094

Thum, T., Caldararu, S., Engel, J., Kern, M., Pallandt, M., Schnur, R., Yu, L., & Zaehle, S. (2019). A new model of the coupled carbon, nitrogen, and phosphorus cycles in the terrestrial biosphere (QUINCY v1.0; revision 1996). Geoscientific Model Development, 12(11), 4781–4802. https://doi.org/10.5194/gmd-12-4781-2019

van der Sleen, P., Vlam, M., Groenendijk, P., Anten, N. P. R., Bongers, F., Bunyavejchewin, S., Hietz, P., Pons, T. L., & Zuidema, P. A. (2015). ^15^N in tree rings as a bio-indicator of changing nitrogen cycling in tropical forests: an evaluation at three sites using two sampling methods. Frontiers in Plant Science, 6, 229. https://doi.org/10.3389/fpls.2015.00229

Van Sundert, K., Radujković, D., & Cools, N. (2020). Towards comparable assessment of the soil nutrient status across scales—Review and development of nutrient metrics. Global Change Biology. https://onlinelibrary.wiley.com/doi/abs/10.1111/gcb.14802

Vicca, S., Stocker, B. D., Reed, S., Wieder, W. R., Bahn, M., Fay, P. A., Janssens, I. A., Lambers, H., Peñuelas, J., Piao, S., Rebel, K. T., Sardans, J., Sigurdsson, B. D., Van Sundert, K., Wang, Y.-P., Zaehle, S., & Ciais, P. (2018). Using research networks to create the comprehensive datasets needed to assess nutrient availability as a key determinant of terrestrial carbon cycling. Environmental Research Letters: ERL [Web Site], 13(12), 125006. https://doi.org/10.1088/1748-9326/aaeae7

Vitousek, P. M., Aber, J. D., Howarth, R. W., Likens, G. E., Matson, P. A., Schindler, D. W., Schlesinger, W. H., & Tilman, D. (1997). Human alteration of the global nitrogen cycle: Sources and consequences. Ecological Applications: A Publication of the Ecological Society of America, 7(3), 737–750. https://doi.org/10.2307/2269431

Walker, A. P., De Kauwe, M. G., Bastos, A., Belmecheri, S., Georgiou, K., Keeling, R. F., McMahon, S. M., Medlyn, B. E., Moore, D. J. P., Norby, R. J., Zaehle, S., Anderson-Teixeira, K. J., Battipaglia, G., Brienen, R. J. W., Cabugao, K. G., Cailleret, M., Campbell, E., Canadell, J. G., Ciais, P., … Zuidema, P. A. (2020). Integrating the evidence for a terrestrial carbon sink caused by increasing atmospheric CO2. The New Phytologist. https://doi.org/10.1111/nph.16866

Zaehle, S., & Dalmonech, D. (2011). Carbon–nitrogen interactions on land at global scales: current understanding in modelling climate biosphere feedbacks. Current Opinion in Environmental Sustainability, 3(5), 311–320. https://doi.org/10.1016/j.cosust.2011.08.008

Zaehle, S., & Friend, A. D. (2010). Carbon and nitrogen cycle dynamics in the O-CN land surface model: 1. Model description, site-scale evaluation, and sensitivity to parameter estimates. Global Biogeochemical Cycles, 24(1), Gb1005. https://doi.org/10.1029/2009GB003521

