## Supplementary tables and figures for "Long-term ecosystem nitrogen limitation from foliar δ^15^N data and a land surface model"

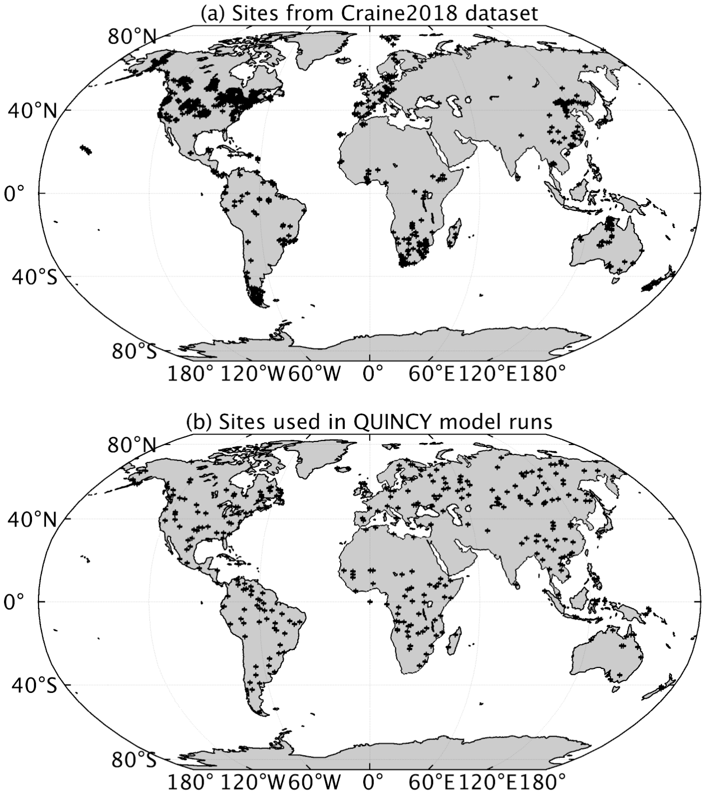


Figure S1 - Geographical distribution of (a) sites from the Craine2018 dataset and (b) sites used for all QUINCY model scenarios.

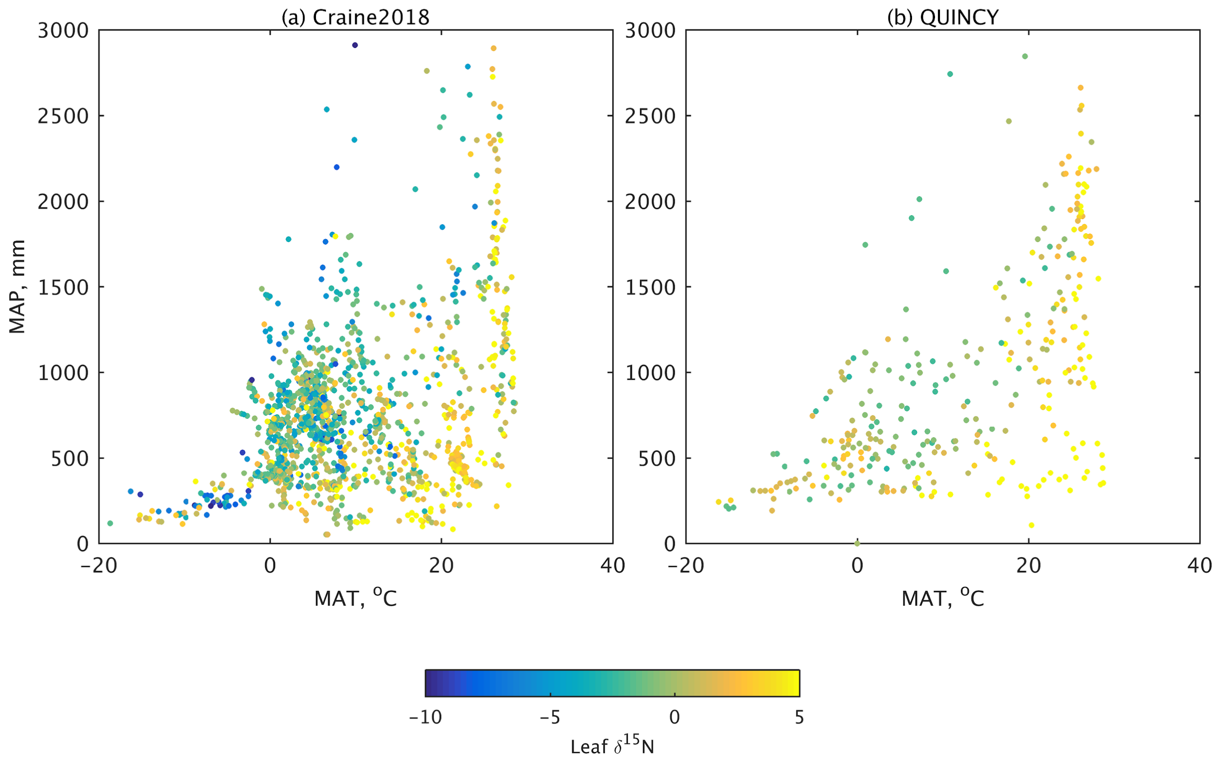


Figure S2 - Distribution of absolute values of leaf $\delta^{15}N$ across climate regions for (a) the Craine 2018 dataset and (b) the QUINCY model.

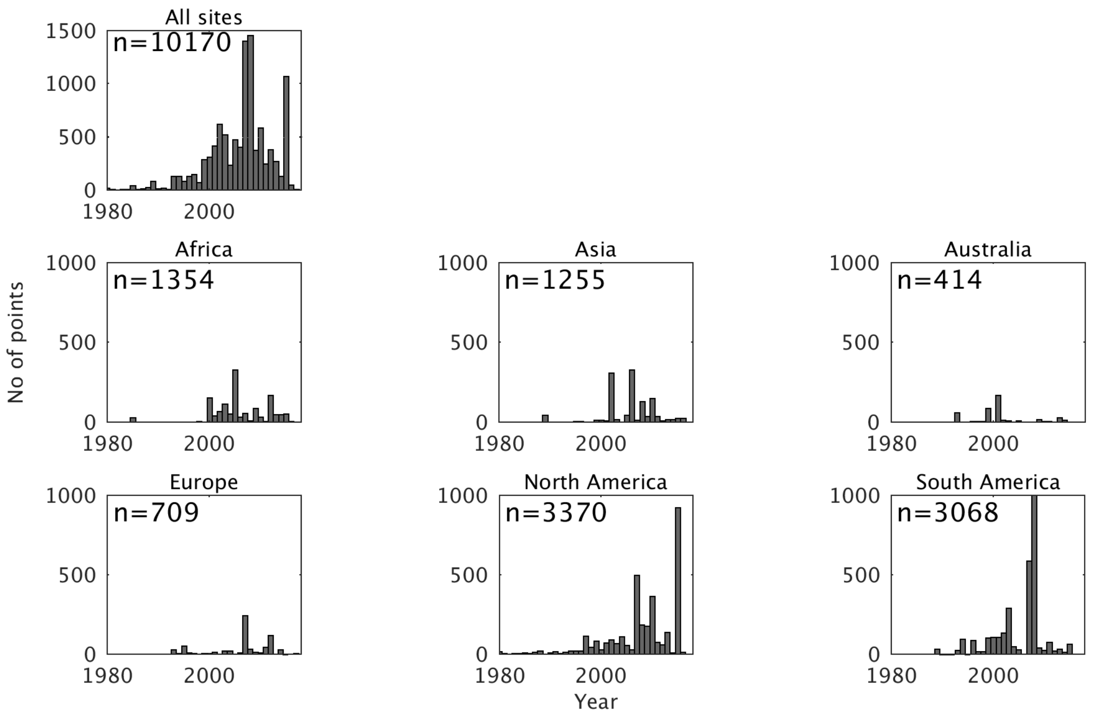


Figure S3 - Temporal distribution of data points in the Craine2018 dataset.

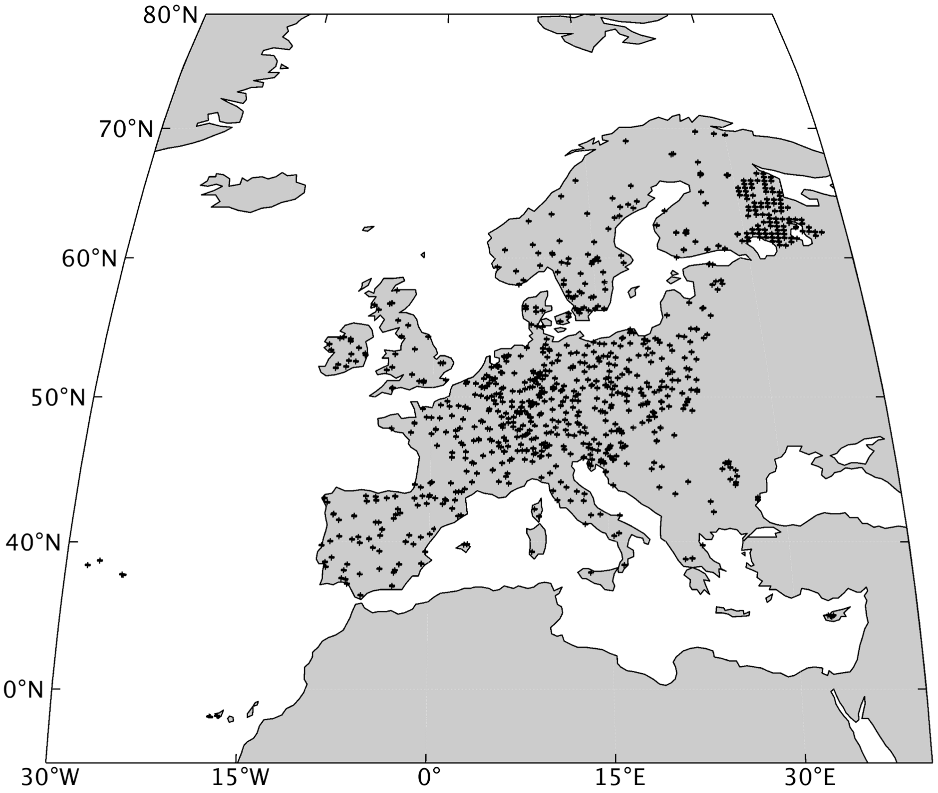


Figure S4 - Geographic distribution of measurement plots in the ICP Forests dataset.

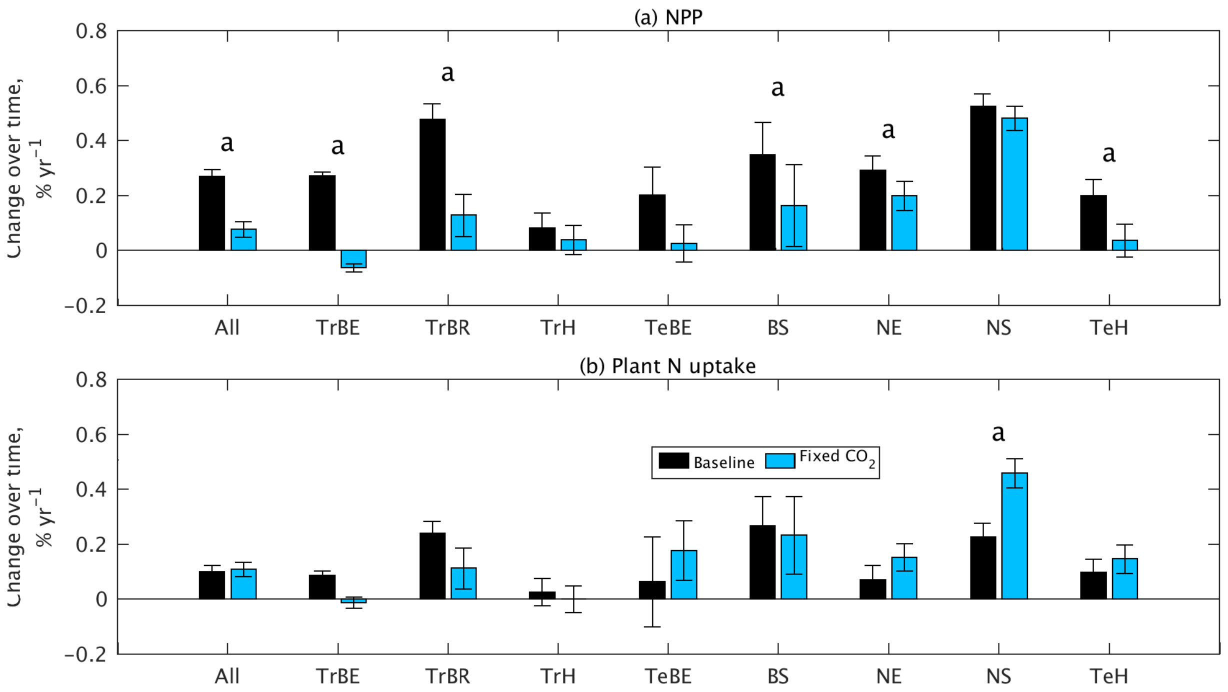


Figure S5 - Plant growth change from 1980-2018 across all PFTs due to the increase in atmospheric CO2 as predicted by the model. Relative change in (a) annual net primary productivity (NPP), (b) plant nitrogen uptake. Error bars represent standard error across sites. PFTs for which the model predicts a significant difference between the baseline and fixed $CO_2$ runs are indicated with ‘a’; significance computed with Wilcoxon rank sum test at $p<0.05$. PFT abbreviations:TrBE tropical broadleaf evergreen,TrBR tropical broadleaf rain-green,TrH, C4 grassland, TeBE temperate broadleaf evergreen, BS broadleaf seasonal, NE needleleaf evergreen, NS needleleaf seasonal, TeH C3 grassland.

Table S1 - Plant functional type (PFT) used in the QUINCY model

| Abbreviation | Description |
|---|---|
| TrBE | Tropical broadleaf evergreen |
| TrBR | (Tropical)broadleaf rain deciduous |
| TeBE | Temperate broadleaf evergreen |
| BS | (Temperate and boreal) broadleaf winter deciduous |
| NE | (Temperate and boreal) needleleaf evergreen |
| NS | (Temperate and boreal) needleleaf winter deciduous |
| TeH | C3 grass |
| TrH | C4 grass |

Table S2 - Nitrogen isotopic fractionation (ε) values in QUINCY, as described in (Thum *et al.*, 2019) adapted from (Robinson, 2001)

| Process | Value, ‰ |
|---|---|
| Microbial $NH_4$ uptake | 17.0 |
| Plant $NH_4$ uptake | 13,5 |
| Plant $NO_4$ uptake | 9.5 |
| Nitrification | 47.5 |
| $NO_3$ production | 25.0 |
| Denitrification | 31.0 |
| $NH_4$ production | 2.5 |
| Biological nitrogen fixation | 3.0 |

Table S3 - Number of sites within each PFT for each dataset used and in the QUINCY model sites

| PFT | QUINCY | Craine2018 | ICP Forests |
|---|---|---|---|
| TrBE | 48 | 3470 | - |
| TrBR | 10 | 1432 | - |
| TeBE | 8 | 364 | 33 |
| BS | 28 | 1492 | 324 |
| NE | 54 | 396 | 613 |
| NS | 9 | 11 | 4 |
| TeH | 74 | 2273 | - |
| TrH | 80 | 743 | - |

Robinson, D. (2001) ‘δ15N as an integrator of the nitrogen cycle’, *Trends in ecology & evolution*, 16(3), pp. 153–162. doi: 10.1016/S0169-5347(00)02098-X.

Thum, T. *et al.* (2019) ‘A new model of the coupled carbon, nitrogen, and phosphorus cycles in the terrestrial biosphere (QUINCY v1.0; revision 1996)’, *Geoscientific Model Development*, 12(11), pp. 4781–4802. doi: 10.5194/gmd-12-4781-2019.
